# Integrated analysis of cervical squamous cell carcinoma cohorts from three continents reveals conserved subtypes of prognostic significance

**DOI:** 10.1101/2020.04.02.019711

**Authors:** Ankur Chakravarthy, Ian Reddin, Stephen Henderson, Cindy Dong, Nerissa Kirkwood, Maxmilan Jeyakumar, Daniela Rothschild Rodriguez, Natalia Gonzalez Martinez, Jacqueline McDermott, Xiaoping Su, Nagayasau Egawa, Christina S Fjeldbo, Vilde Eide Skingen, Mari Kyllesø Halle, Camilla Krakstad, Afschin Soleiman, Susanne Sprung, Peter Ellis, Mark Wass, Martin Michaelis, Heidi Lyng, Heidi Fiegl, Helga Salvesen, Gareth Thomas, John Doorbar, Kerry Chester, Andrew Feber, Tim R Fenton

## Abstract

Human papillomavirus (HPV)-associated cervical cancer represents one of the leading causes of cancer death worldwide. Although low-middle income countries are disproportionately affected, our knowledge of the disease predominantly originates from populations in high-income countries. Using the largest multi-omic analysis of cervical squamous cell carcinoma (CSCC) to date, totalling 643 tumours and representing patient populations from the USA, Europe and Sub-Saharan Africa, we identify two CSCC subtypes (C1 and C2) with differing prognosis. C1 tumours are largely HPV16-driven, display increased cytotoxic T-lymphocyte infiltration and frequently harbour *PIK3CA* and *EP300* mutations. C2 tumours are associated with shorter overall survival, are frequently driven by HPVs from the HPV18-containing alpha-7 clade, harbour alterations in the Hippo signalling pathway and increased expression of immune checkpoint genes, *B7-H3* (also known as *CD276*) and *NT5E* (also known as *CD73*) and *PD-L2* (also known as *PDCD1LG2*). In conclusion, we identify two novel, therapy-relevant CSCC subtypes that share the same defining characteristics across three geographically diverse cohorts.

---

Despite screening and the introduction of prophylactic human papillomavirus (HPV) vaccination in developed countries, cervical cancer continues to be one of the leading worldwide causes of cancer-related deaths in women^1^. Prognosis for patients with metastatic disease remains poor, thus new treatments and effective molecular markers for patient stratification are urgently required. Cervical cancer is caused by at least 14 high-risk human papillomaviruses (hrHPVs), with HPV16 and HPV18 together accounting for over 70% of cases worldwide, with some variation by region^1,2^. Cervical squamous cell carcinoma (CSCC) is the most common histological subtype of cervical cancer, accounting for approximately 60-70% of cases, again with some variation seen across different populations^2^. Adeno- and adenosquamous histology are both associated with poor prognosis^3–6^, while the relationship, if any, between HPV type and cervical cancer prognosis remains unclear^7^. HPV type is also associated with histology, with HPV16 more commonly found in CSCC, while adenocarcinomas are more likely to harbour HPV18^2^. Previous landmark studies described the genomic landscape of cervical cancer in different populations^8–11^ and in some cases identified subtypes based on gene expression, DNA methylation and/or proteomic profiles^8,9^. The Cancer Genome Atlas (TCGA) network identified clusters based on RNA, micro-RNA, protein/phospho-protein, DNA copy number alterations and DNA methylation patterns and combined data from multiple platforms to define integrated iClusters^8^. In their analysis, only clustering based on the expression levels and/or phosphorylation state of 192 proteins as measured by reverse-phase protein array (RPPA) was associated with outcome, with significantly shorter overall survival (OS) observed for a cluster of cervical cancers exhibiting increased expression of Yes-associated protein (YAP) and features associated with epithelial-to-mesenchymal transition (EMT) and a reactive tumour stroma. Since TCGA’s RPPA analysis was restricted to 155 tumours including SCCs, adeno- and adenosquamous carcinomas, we set out to test the hypothesis that with data from more samples, we could identify a set of transcriptional and epigenetic features associated with prognosis within CSCC and to establish whether it is also present in independent patient cohorts representing different geographical locations and ethnicities. To identify molecular subtypes and prognostic correlates, we identified a set of 643 CSCCs (all HPV-positive), for which clinico-pathological data and genome-wide DNA methylation profiles were either publicly available or generated in this study, and for which in most cases, matched gene expression and somatic mutation data were also available (Table 1).

**Table 1:**
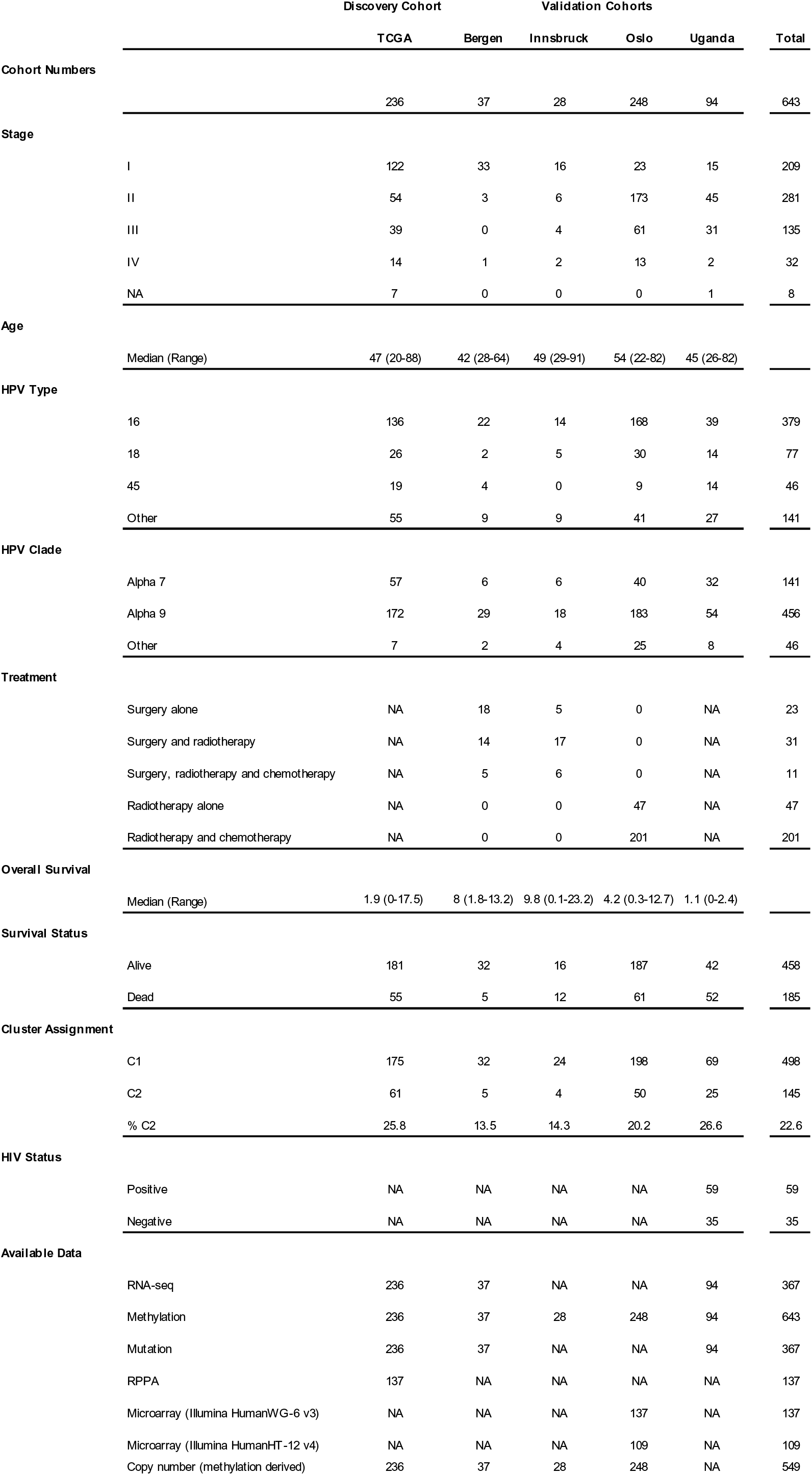
Summary of clinicopathological characteristics for five cervical cancer cohorts.

## Identification of two gene expression-based clusters in cervical squamous cell carcinoma

Molecular and clinical differences between cervical adeno/adenosquamous and CSCCs are well documented^12–14^ and gene expression differences were apparent in multi-dimensional TSNE analysis based on the top 10% most variable genes of three previously published cervical cancer cohorts^8,9,11^ with available RNA-seq data (Supplementary Fig. S1a-d). To examine molecular and clinical heterogeneity specifically within SCC we focused all subsequent analysis on a collection of confirmed HPV-positive CSCCs from the USA, Europe and Uganda, as shown in Table 1.

236 cervical SCCs profiled by TCGA were defined as our discovery cohort (Table 1, Supplementary Table S1) and consensus clustering was performed using the top 10% most variable genes (n=1377 genes, Supplementary Table S2). Consensus cluster membership heatmaps, delta area plot, consensus cumulative distribution function (CDF) and proportion of ambiguous clusters (PAC) indicated the optimal number of clusters was two (Fig. 1a, Supplementary Fig. 2), the larger of which (n=175) was designated C1 while the smaller cluster (n=61) was designated C2 (Supplementary Table S1). Modelling transcriptomic differences between these two clusters identified 938 differentially expressed genes (DEGs, FDR=0.01, FC > 2) (Fig. 1b, Supplementary Table S3). Tumours in C1 predominantly harbour HPV types from the HPV16-containing alpha-9 clade (150/175) while 38 of 61 C2 tumours contained HPV types from the HPV18-containing alpha-7 clade. C2 tumours were 13.3 times more likely to harbour alpha 7 HPVs than C1 tumours (p = 1.8 x 10^-14^, Fishers Exact Test) (Fig. 1b).

**Figure 1.**
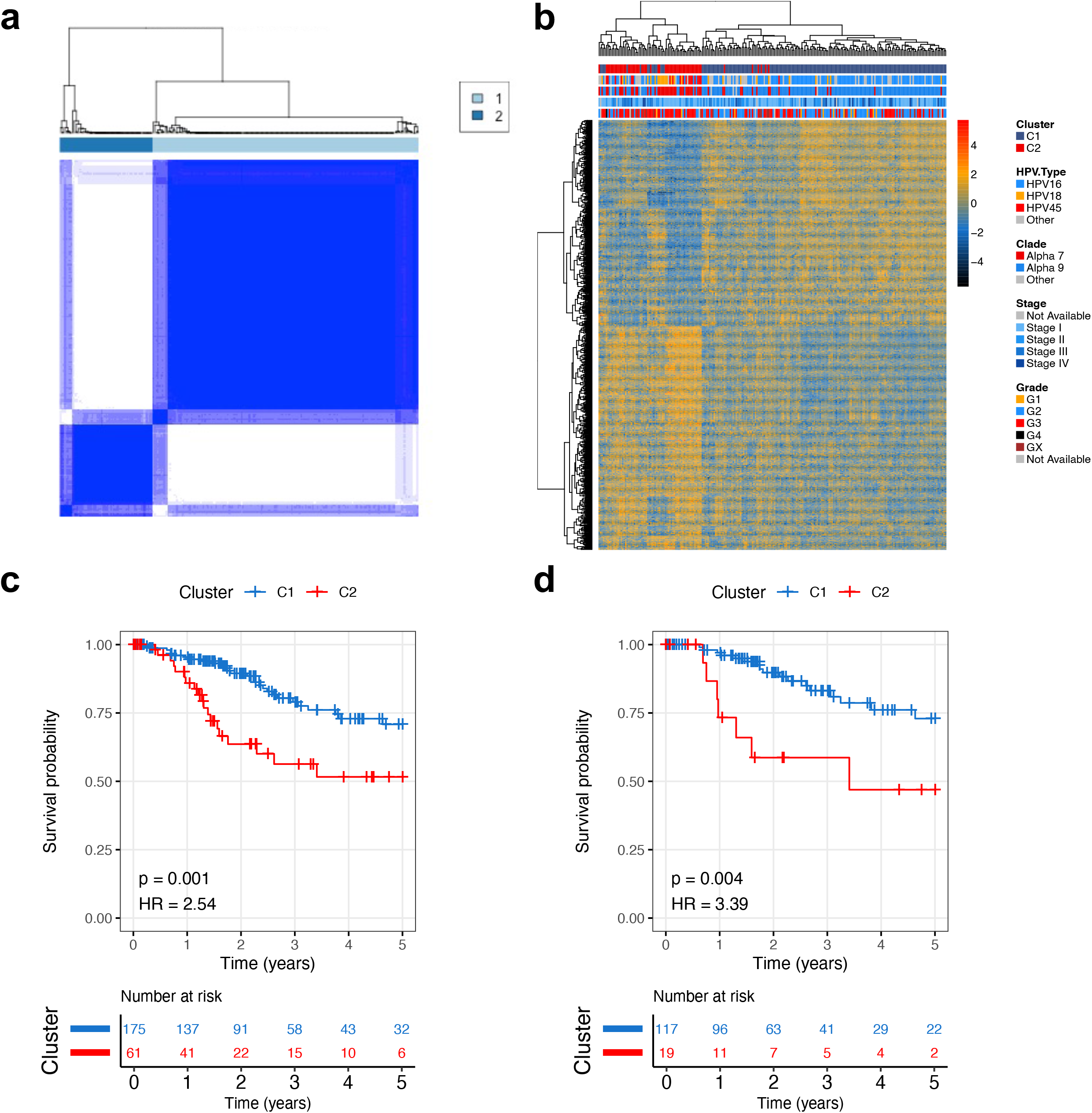
Consensus clustering produces two prognostic clusters in TCGA SCC cohort. **a)** Consensus clustering of 236 TCGA HPV+ SCC patients. **b)** There were 938 differentially expressed genes between the two clusters. **c)** 5 year survival between the 2 SCC subgroups. **d)** 5 year survival between the 2 SCC subgroups considering only HPV16+ tumours. Statistics from univariate Cox regression.

Univariate analysis of 5-year overall survival (OS) revealed worse outcomes for patients with C2 tumours (HR = 2.54, p = 0.001; Fig. 1c) and in Cox regression including age, tumour stage and HPV type as covariates along with cluster membership, only membership of the C2 cluster (HR = 2.44, p = 0.017 95% CI 1.18, 5.05) and a tumour stage of IV (HR versus stage I = 4.74, p<0.001, 95% CI 2.1, 10.7) were independent predictors of five-year OS (Table 2). The relationship between cluster and OS is also clear when restricting the analysis to HPV16-containing tumours in each cluster in both univariate analysis (HR = 3.39, p = 0.004; Fig. 1d) and multivariate analysis, including age and tumour stage as covariates (HR = 3.89, p = 0.003, 95% CI 1.57, 9.67; Supplementary Table S4).

**Table 2.** Five-year survival analysis for all cohorts

|  | Univariate |  |  | Multivariate |  |  |
| --- | --- | --- | --- | --- | --- | --- |
|  | Hazard Ratio | p Value | 95% CI | Hazard Ratio | p Value | 95% CI |
| **TCGA** | 2.54 | 0.002 | 1.42, 4.56 | 2.44 | 0.02 | 1.18, 5.05 |
| **Bergen** | 5.28 | 0.07 | 0.87, 31.9 | 98.1 | <0.001 | 8.41, 1145 |
| **Innsbruck** | 0 | 1 | 0, Inf | 0 | 1 | 0, Inf |
| **Oslo** | 1.74 | 0.07 | 0.96, 3.14 | 2.36 | 0.012 | 1.21, 4.62 |
| **Europe Combined** | 1.68 | 0.07 | 0.97, 2.90 | 2.54 | 0.003 | 1.40, 4.67 |
| **Uganda** | NA | NA | NA | NA | NA | NA |

## Identification of C1 and C2 CSCCs and association with prognosis in independent SCC cohorts

To further investigate the association between C1/C2 cluster membership and OS, we assembled a combined validation cohort consisting of 313 CSCC patients treated at three centres in Europe (Bergen (n = 37), Oslo (n = 248) and Innsbruck (n = 28)), for which detailed clinical information were available and for which genome-wide DNA methylation profiles from Illumina Infinium 450k arrays (the same platform used by TCGA) were either available or generated in this study (Table 1). Since RNA-seq data were not available for all European samples, cluster membership was assigned using a support vector machine (SVM) classification model based on 129 CpG sites (methylation variable positions, MVPs) at which methylation differed significantly between tumours in C1 vs C2 clusters in the discovery cohort (Fig. 2a, b; mean delta-Beta > 0.25, FDR < 0.01, Supplementary Table S5), 18 of which were located within 12 genes differentially expressed between the clusters (Supplementary Fig. S3). MVP and DEG signatures were also used to assign cluster membership to 94 CSCCs from a Ugandan cohort originally profiled by the Cancer Genome Characterization Initiative (CGCI)^9^, for which both DNA methylation and RNA-seq data were available. C2 tumours from all cohorts clustered together using TSNE analysis based on the MVP signature (Fig. 2c) and high concordance between DEG and MVP-based cluster allocation was observed in all cohorts for which both gene expression (RNA-seq for Uganda and Bergen or Illumina bead chip arrays for Oslo) and DNA methylation data were available (Supplementary Fig. S4a, b). Single-sample gene set enrichment analysis (ssGSEA) confirmed differential expression of the signature genes in tumours classified as C1 or C2 using DNA methylation data (Supplementary Fig. 4c). 59 of 313 (18.8%) tumours in the combined European cohort (Fig. 2b, Supplementary Table S7) and 25 of 94 (26.6%) tumours in the Ugandan cohort were classified as C2 (Supplementary Fig. S5a, Supplementary Table S6). As in the discovery cohort, most C1 tumours from the European and Ugandan cohorts harboured alpha-9 HPV types (260/325) while C2 tumours were 3.9 times more likely to harbour alpha-7 HPVs than C1 tumours (p = 1.07 x 10^-6^, Fishers Exact Test) (Fig. 2b, Supplementary Fig. S5a). Interestingly 80% (20/25) of Ugandan C2 patients were human immunodeficiency virus (HIV) positive, while only 56% (39/69) of C1 patients were HIV positive (Supplementary Fig. S5a).

**Figure 2.**
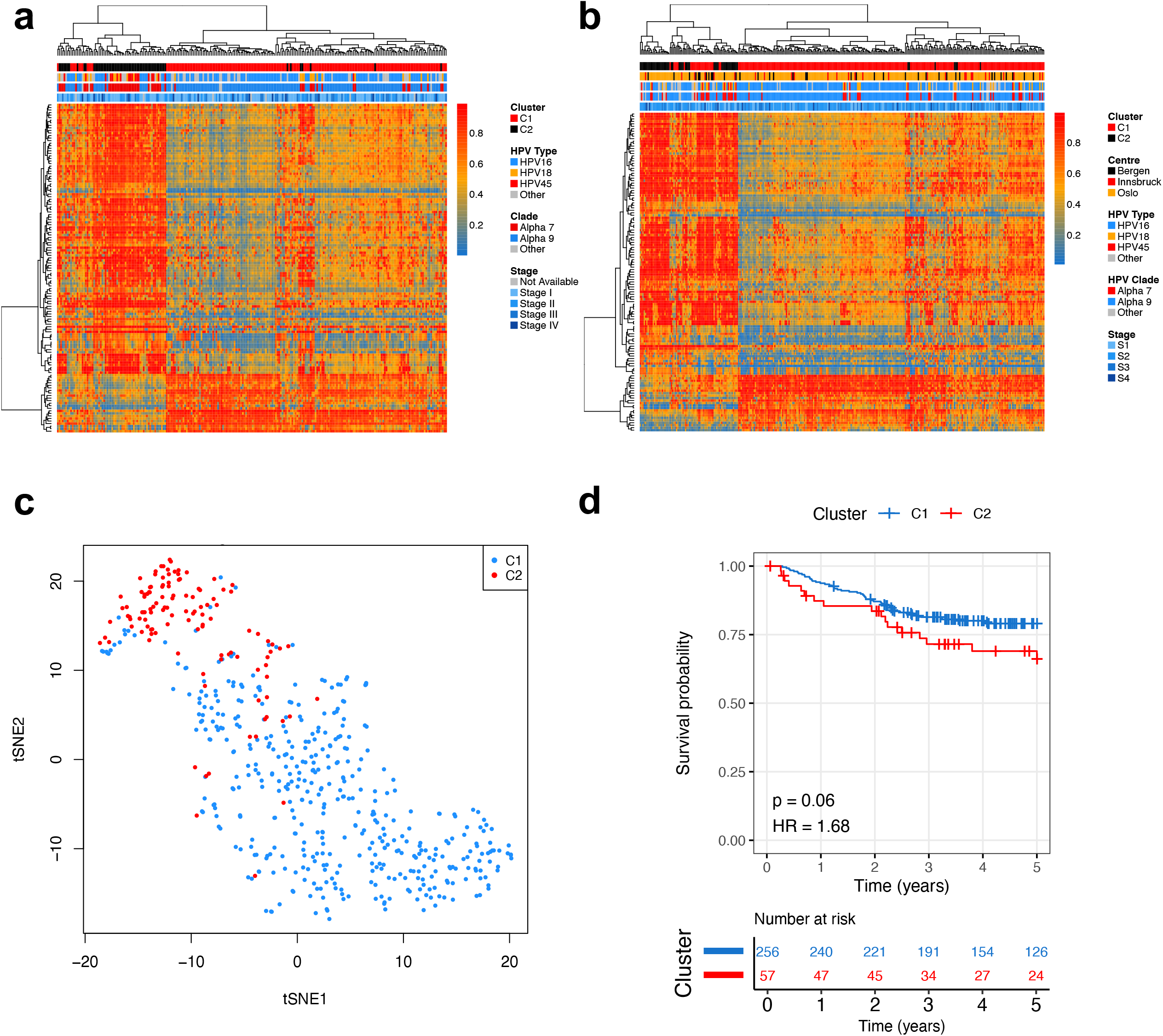
Cluster allocation of validation cohorts using methylation signature. **a)** A signature of DNA methylation (dB > 0.25, FDR < 0.01) separates C1 and C2 SCC subgroups in the TCGA cohort. **b)** The methylation patterns are reproduced in a validation dataset from three European centres (n = 313). **c)** C2 tumours from TCGA and European validation cohorts cluster together based on the 129 MVP signature. **d)** 5 year survival curve for combined European validation cohorts. Statistics from univariate Cox regression.

Univariate analysis indicated lower 5-year OS in C2 tumours from the European cohort (Fig. 2d) and Cox regression controlling for FIGO stage, age, HPV type and treatment (surgery alone, surgery with radio-chemotherapy, surgery with radiotherapy alone, radio-chemotherapy and radiotherapy alone) again identified C2 status but not HPV type to be an independent predictor of 5-year OS (HR = 2.54, p =0.003, 95% CI 1.4, 4.7) along with tumour stage and inclusion of chemotherapy in the treatment regimen (Table 2). As in the discovery cohort, a significant prognostic difference was identified between the C1 and C2 subgroups when considering only the HPV16-positive tumours (n = 204) in both univariate (Supplementary Fig. S5b) and multivariate analyses (HR = 2.64, p = 0.02, 95% CI = 1.16, 6; Supplementary Table S4). Interestingly the prognostic difference was even greater among 78 patients in the European cohort that did not receive chemotherapy (Supplementary Fig. S5c; multivariate HR = 4.4, p = 0.005, 95% CI = 1.58, 12.3). At 94 patients, the Ugandan cohort was underpowered for comparing survival between C1 and C2 tumours and survival rates in the Ugandan cohort were much lower than in the other cohorts (Supplementary Fig. S5d), thus we did not attempt a combined survival analysis including these patients. Taken together, the C1/C2 clusters identified in the TCGA cohort (USA) are apparent in cohorts of CSCC patients from Europe and Uganda and tumours can be accurately assigned to cluster using either gene expression or DNA methylation profiles. C1/C2 cluster is an independent predictor of 5-year OS in both the TCGA (n = 236) and European (n = 313) cohorts and remains so when only HPV16+ tumours are considered. There is no difference in the breakdown of C1 and C2 tumours by stage (Supplementary Table S7).

## Relationships between C1/C2 and clusters previously identified by TCGA

Of the 178 tumour samples that made up the core set in the TCGA’s landmark study into cervical cancer genomics/epigenomics^8^, 140 CSCCs were present in our discovery cohort of 236 (Supplementary Table S8). This enabled comparisons between our gene expression-based cluster allocations and the subtypes defined by TCGA (Fig. 3). TCGA analysis included integrated clustering using multiomics data (three iClusters, two of which (‘keratin-high’ and ‘keratin-low’ were composed entirely of CSCCs) and clustering based on transcriptomic data (three mRNA clusters). There is considerable overlap between our C1 cluster and TCGA’s mRNA C2 cluster (84/106) and keratin-high iCluster (80/106), and between our C2 cluster and TCGA’s mRNA C3 cluster (19/34) and keratin-low iCluster (27/34). Neither the mRNA C3 nor the keratin-low iCluster were associated with poor prognosis in TCGA’s analysis and given the increased expression of a subset of keratin genes (including *KRT7*, *KRT8* and *KRT18*) in C2 tumours (Fig. 3), we decided against adopting the keratin-high / keratin-low nomenclature for our clusters. We also examined the relationship between our subtypes and three clusters defined by TCGA based on reverse phase protein array (RPPA) data. Notably, 57% of C2 TCGA tumours with RPPA data available belong to the EMT cluster compared with only 25% of C1 tumours (Fig. 3) and, consistent with the proteomic classification, C2 tumours display higher EMT mRNA expression scores, as defined by TCGA^8^ than C1 tumours (Supplementary Fig. S6). Although there is greater concordance between C2 and the TCGA EMT cluster compared to C1, it is clearly distinct from the EMT cluster.

**Figure 3.**
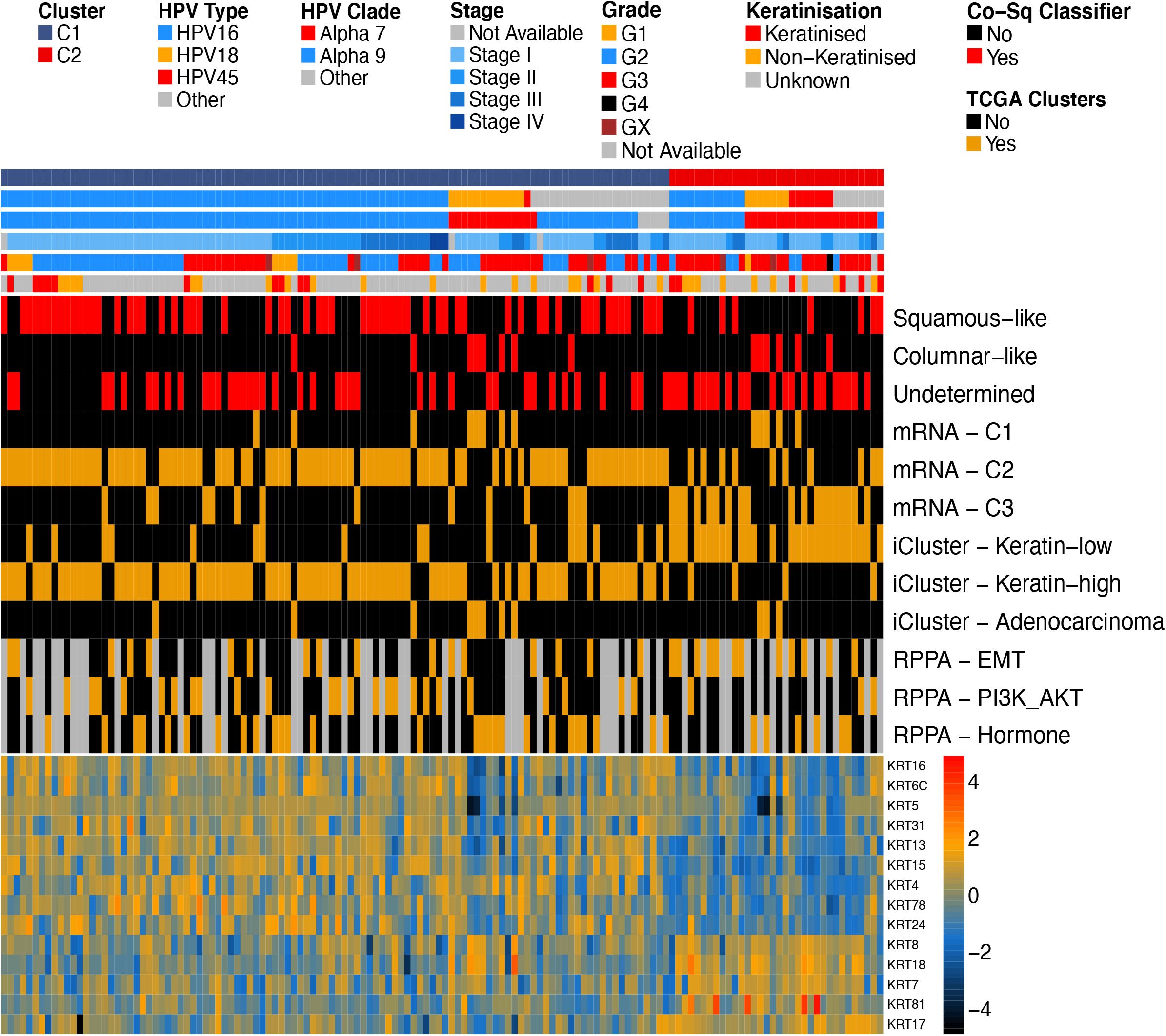
Comparison of SCC subgroups with previous studies. Cluster analysis had previously been performed on 140 TCGA SCC tumours in two studies – one determined clusters based on cell of origin markers (Chumduri *et al,* 2021, red), one determined clusters based on integrated omics data (TCGA Network, 2017, orange). The heatmap at the bottom of plot represents expression levels of cytokeratin genes present in our C2 gene signature.

## Genomic analyses of prognostic clusters

To investigate whether C1 and C2 tumours differ at the genomic level in addition to the transcriptomic and epigenomic differences observed above, whole-exome data was obtained for SCCs from three cohorts, TCGA^8^, Bergen^11^ and Uganda^9^. This amounted to 367 samples, 29 of which were classed as hypermutated by standards set by TCGA^8^ (>600 mutations). The median tumour mutation burden (TMB) was 2.04/Mb for all tumour, 2.11/Mb for C1 tumours and 1.82/Mb for C2 tumours (1.92/Mb, 1.94/Mb and 1.72/Mb respectively after removal of hypermutated samples). We detected four mutation signatures for the combined cohorts (Supplementary Fig. S7): as expected based on previous studies^8,9,11^, COSMIC signatures 2 and 13 (characterised by C>T transitions or C>G transversions respectively at TpC sites attributed to cytosine deamination by APOBEC enzymes); age-related COSMIC signature 1 (characterised by C>T transitions attributed to spontaneous deamination of 5’ methylated cytosine) and COSMIC signature 5, for which the underlying mutational process is unknown^15^ (https://cancer.sanger.ac.uk/signatures/). The proportion of mutations attributable to each signature did not vary between clusters (Fig. 4).

**Figure 4.**
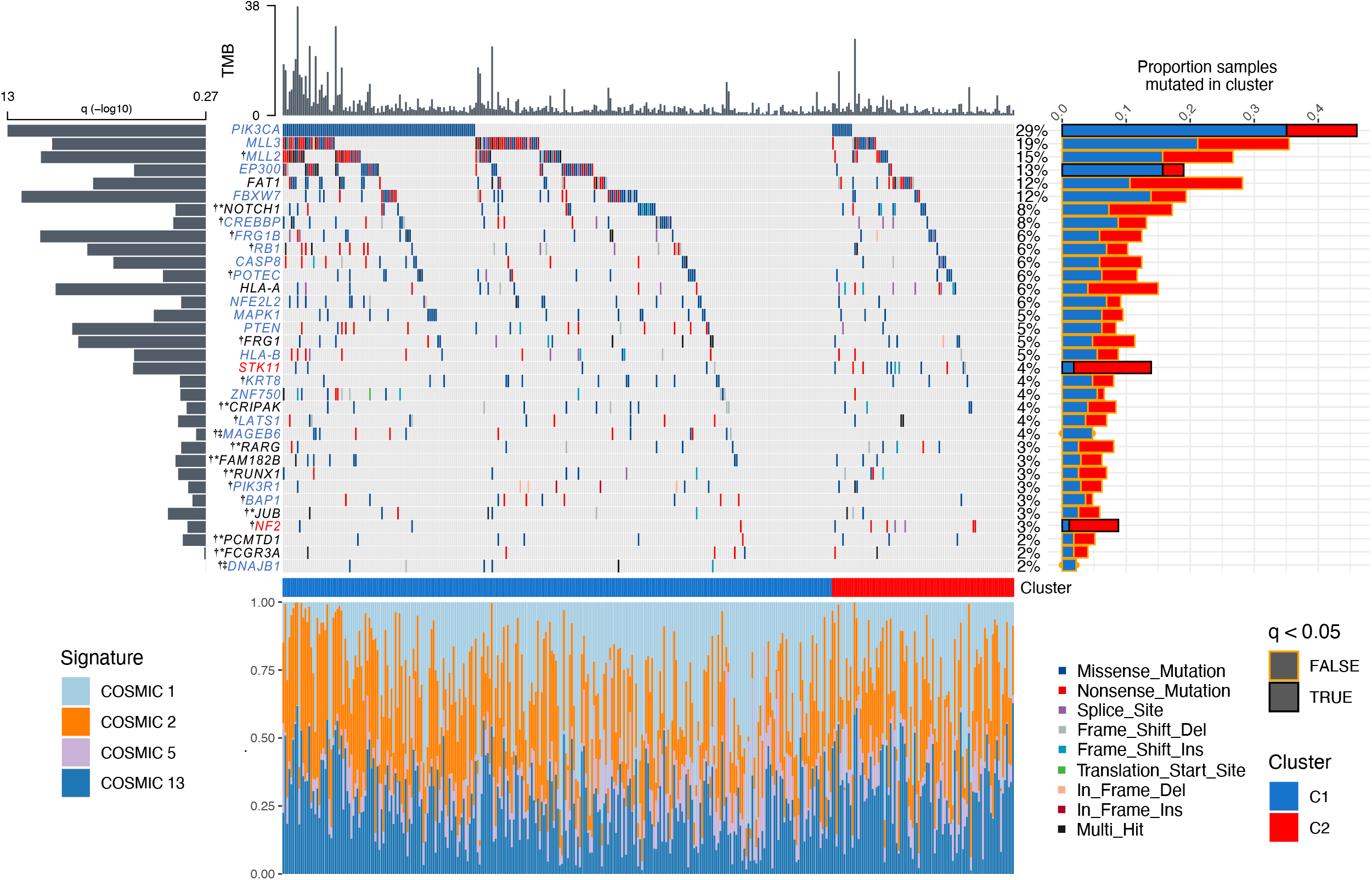
Genomic summary of significantly mutated genes (SMGs) in SCC cohorts. Main plot shows mutation type and frequencies for 34 SMGs identified using dNdSCV on TCGA, Bergen and Ugandan cohorts (367 total patients). Grey bars at top of plot represent TMB per sample. Grey bars to left of plot represent significance of SMG, larger bar is more significant. Barchart to the right shows proportion of a genes mutations in by cluster (blue = C1, red = C2). Black box around bar represents a significant difference in mutation frequency between the clusters (p<0.05) while a gold box means no significant difference between the clusters. The plot at the bottom of figure represents the mutational signatures that contribute towards each individuals tumour mutational burden. [Gene name key – blue – unique to C1 analysis, red = unique to C2 analysis, black = both in C1 and C2 individual analyses, black* = only significant when combining both clusters for analysis, **†** = novel SMG in cervical cancer, **‡** = not significant in combined cluster analysis but significant in C1 only analysis].

Having excluded the hypermutated samples, we next performed dNdScv analysis^16^ on each cohort, followed by p-value combination using sample size weighted Fisher’s method followed by FDR correction^17^ to permit identification of significantly mutated genes (SMGs) across the entire dataset. This combined approach, followed by analysis of individual samples by cluster identified 34 SMGs (Fig. 4, Supplementary Table S9), 21 of which (highlighted by **^†^**) have not previously been identified as SMGs in cervical cancer^8,9,11^. Of the 34 SMGs, 21 were significantly mutated in only C1 samples, two genes in only C2 samples, three genes in both C1 and C2 individual analysis, and eight genes were only significantly mutated when both C1 and C2 clusters were analysed together (Fig. 4, Supplementary Table S9). The frequency of mutations in SMGs that had been previously observed was comparable between combined cohort and each respective SMG study (Supplementary Table S10). Among the 21 genes that have not previously been identified as significantly mutated in cervical cancer, six are SMGs in other SCCs, including head and neck (*NOTCH1, JUB (*also known as *AJUBA), MLL2 (*also known as *KMT2D), RB1, PIK3R1*)^18^, oesophageal (*MLL2, NOTCH1, RB1*)^19^ and lung SCC (*NOTCH1, RB1, MLL2, CREBBP (*also known as *KAT3A*))^20^. Conversely, several genes previously identified as SMGs in cervical cancer, including *TP53*, *ARID1A* and *TGFBR2* are significantly mutated in adenocarcinoma but not in CSCC^8,11^. Comparing somatic mutation rates in SMGs between clusters using binomial regression identified *PIK3CA* (FDR = 0.001) and *EP300* (FDR = 0.046) mutations as disproportionally more common in C1 tumours and *STK11* (FDR = 0.005) and *NF2* (FDR = 0.045) as enriched in C2 tumours (Fig. 4). *STK11* is also under-expressed in C2 tumours compared with C1 tumours (Supplementary Table S3).

## C2 tumours display Hippo pathway alterations and increased YAP1 activity

Two SMGs from our analysis (*LATS1* and *NF2*) are core members of the HIPPO signalling pathway, while SMGs *FAT1, JUB* and *STK11* are known regulators of HIPPO signalling^21–23^. Mutations in *LATS1*, *FAT1, JUB, STK11* or *NF2* (the latter two of which are significantly mutated specifically in C2 tumours, Fig. 4) result in aberrant activation of the downstream transcription factor, yes1 associated transcriptional regulator (YAP1)^24–28^, the expression of which is also elevated at the mRNA level in C2 tumours (Table S3).

We generated segmented copy number data for all tumours (combining TCGA and European validation cohort samples for which the necessary data were available for maximum statistical power), which identified 211 focal candidate copy number alterations (CNAs) at FDR < 0.1. Following binomial regression, we identified five discrete CNAs that differed in frequency between C1 and C2 clusters (Fig. 5a; FDR < 0.1, log2 (Odds Ratio) > 1). All five were more prevalent in C2 tumours and included 11q11 and 1q21.2 deletions and 6p22.1, 11q22.1 and 11q22.2 gains. 11q22.2 contains matrix metalloproteinase genes (MMPs) which are well known to be involved in metastasis^29^, but notably 11q22.1 contains the *YAP1* gene. Furthermore, analysis of Reverse Phase Protein Assay (RPPA) data from TCGA revealed significantly higher YAP1 protein expression in C2 tumours (Fig. 5b). We confirmed that of the 137 TCGA cases for which RPPA data were available, cases with *YAP1* amplification (8/37 C2 tumours and 6/100 C1 tumours) also showed increased YAP1 mRNA and protein expression (Supplementary Fig. S8). In total 10 genes from a 22 gene signature that predicts HIPPO pathway activity in cancer^30^ are differentially expressed between C1 and C2 tumours (Supplementary Table S3).

**Figure 5.**
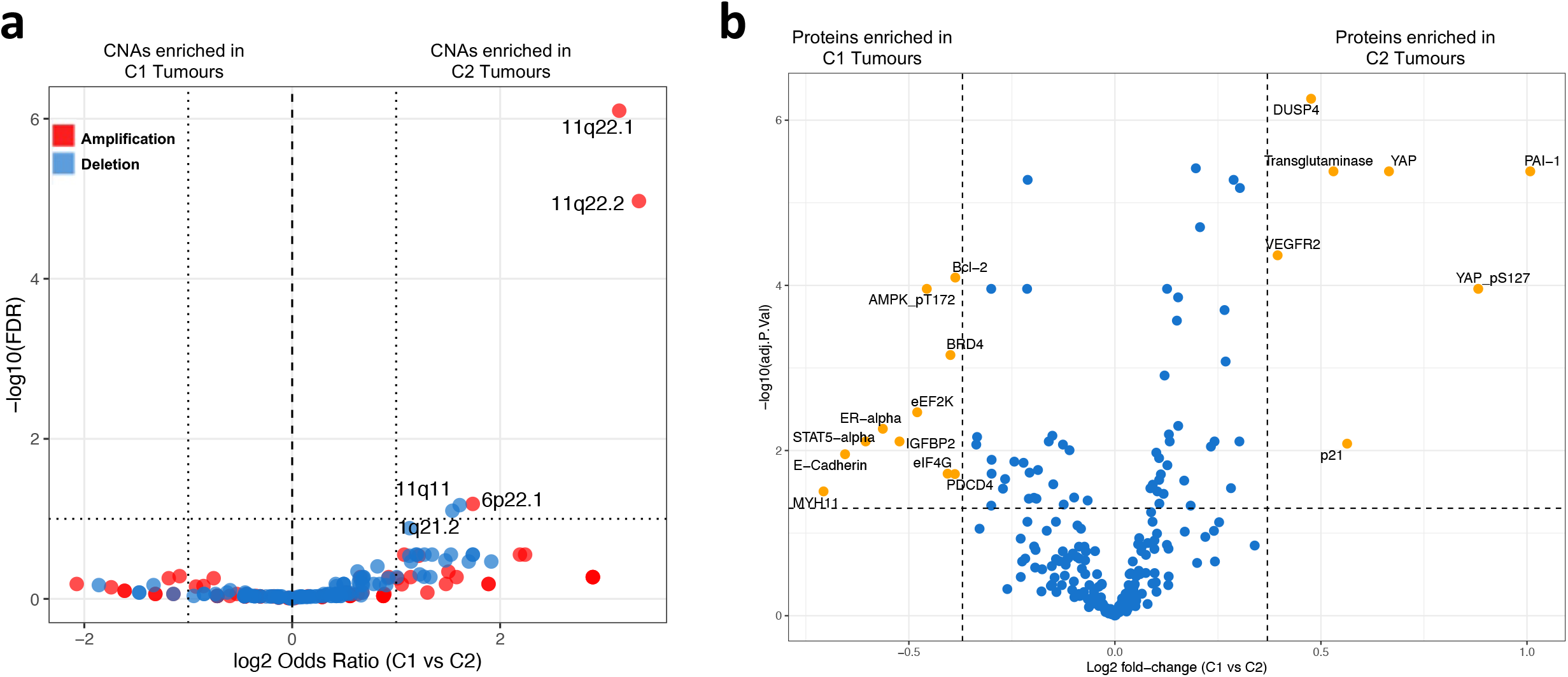
Copy number and protein level differences between SCC subgroups. **a)** Volcano plot showing differences in GISTIC copy number peak frequencies between C1 and C2 tumours, with –log10(FDR) on the y axis and the odds ratio on the x axis. **b)** Volcano plot showing differentially abundant proteins and phospho-proteins (FDR < 0.05, FC > 1.3, represented by yellow dots) between C1 and C2 TCGA tumours, as measured by Reverse Phase Protein Array.

## Differences in the tumour immune microenvironment between C1 and C2 tumours

The nature of the tumour immune microenvironment, particularly the abundance of tumour infiltrating lymphocytes (TILs) is a strong prognostic factor in cervical cancer^31–33^. We used DNA methylation data to compare the cellular composition of TCGA tumours^34^, observing differences in the proportions of multiple cell types between the subgroups (Fig. 6a); most notably decreased CD8+ (cytotoxic T lymphocytes (CTL)), and a marked elevation of neutrophil and CD56+ natural killer (NK)-cells in C2 tumours. Repeating this method with the validation cohorts produced results that were remarkably similar (Fig. 6b). Differences in the proportions of cell types between C1 and C2 in the validation cohort mirrored those in the TCGA cohort, decreased CTL, and elevated neutrophil, NK-cell and endothelial cell levels were observed in C2 tumours. Importantly, this was not driven by any single validation cohort, as individual cohorts displayed consistent patterns of differences in the proportion of cell types between C1 and C2 tumours, especially with regards to CTLs, neutrophils and NK-cells (Supplementary Fig. S9a-d). C2 tumours also exhibit markedly higher neutrophil:CTL ratios (Supplementary Fig. S9e, f) and neutrophil:lymphocyte (CTL, B-cell and Treg) ratios (NLR, Supplementary Fig. S10); established adverse prognostic factors in cervical cancer^35–37^. At 0.7, the NLR in C1 tumours across all cohorts was less than half that observed in C2 tumours (1.85).

**Figure 6.**
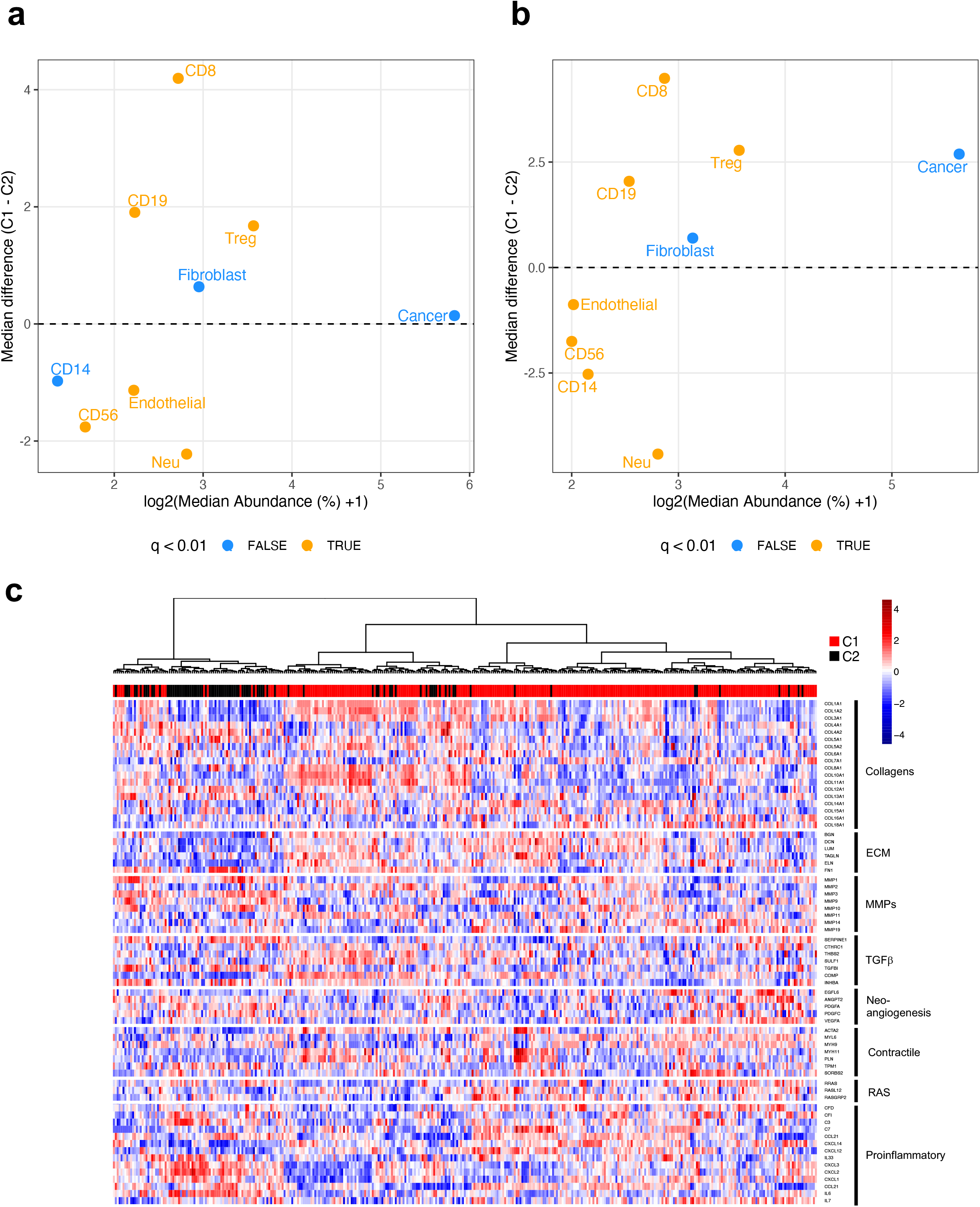
Differences in the tumour microenvironment between cervical cancer subgroups. Plot showing median abundances (x-axis) and median differences (%, y-axis) for different cell types estimated using MethylCIBERSORT, with significant differences in orange, for **a)** TCGA discovery cohort and **b)** combined validation cohorts. **c)** C2 tumours cluster together using CAF geneset genes.

Validation of MethylCIBERSORT cell estimates was performed for a subset of samples from the Innsbruck cohort using CD8 (CTLs) and myeloperoxidase (MPO, neutrophils) immunohistochemistry (IHC)-based scores from a pathologist blinded to cluster designation (Supplementary Fig. S11a-c) and for CTLs in the Oslo cohort samples using comparison of MethylCIBERSORT estimates to CD8 IHC-based digital pathology scores (Supplementary Fig. S11d).

Also of potential significance regarding the tumour immune microenvironment, is the presence of two immune checkpoint genes, *CD276* (also known as *B7-H3*) and NT5E (also known as *CD73*) in the set of 938 signature DEGs that separate the clusters (Table S3). Both *B7-H3* and *NT5E*, along with a third immune checkpoint gene (*PD-L2*) are expressed at higher levels in C2 tumours (Supplementary Fig. S12) and hypomethylation of two CpGs in the *NT5E* promoter is evident in C2 tumours (Supplementary Table S5). All three suppress T-cell activity^38–40^ and *B7-H3* expression has been linked to poor prognosis in cervical cancer^41,42^.

## Evidence for differences in stromal fibroblast phenotype between C1 and C2 tumours

Gene set enrichment analysis using Metascape^43^ suggested increased EMT (Supplementary Table S9) in C2 tumours, with 52 of 200 genes in in the EMT Hallmark gene set upregulated. As noted above there is also greater overlap between the C2 cluster and an EMT cluster defined by TCGA and based on RPPA data (Figure 3). Single-cell RNA sequencing and xenografting studies strongly suggest that rather than arising from the tumour cells (few of which have undergone EMT at any given time^44–46^), mesenchymal gene signatures in bulk tumour expression data instead derive from stromal cells including fibroblasts, which can adopt various phenotypes and play an important role in shaping the tumour immune microenvironment^47,48^. In addition to YAP1, which has been linked to the formation of cancer-associated fibroblasts (CAFs)^49^ (as well as EMT^50–52^ and angiogenesis^53^), C2 tumours display increased expression of the CAF marker genes *FAP* and *SERPINE1* (also known as *PAI-1*)^54^; the latter evidenced at both mRNA and protein levels (Supplementary Table S3, Fig. 5b). Overall fibroblast content as estimated by MethylCIBERSORT is similar between C1 and C2 tumours (Fig. 6a, b) but given recent findings regarding the extent and prognostic significance of CAF heterogeneity in the tumour microenvironment^55–59^, we hypothesized that CAF phenotype rather than overall abundance, may differ between C1 and C2 tumours. To examine this, hierarchical clustering was performed based on the expression of eight gene sets (68 genes) curated by Qian et al^56^, representing CAF-related biological processes and which are differentially expressed across six CAF phenotypes recently identified in a pan-cancer analysis^59^. C2 tumours cluster together, displaying increased expression of proinflammatory genes associated with an inflammatory (pan-iCAF2) CAF phenotype, C1 tumours appear more heterogenous with respect to expression of the signature genes used to define CAF phenotypes; there is upregulation of assorted myofibroblastic (myoCAF) genes in a subgroup C1 tumours, including various collagens, ECM genes and TGFb-associated genes, as well as ‘contractile’ genes such as smooth muscle actin (*ACTA2*, Fig. 6c). While *ACTA2* is commonly used to identify myoCAF, it is also expressed by pericytes and smooth muscle cells, which share the contractile phenotype (and express for example, *MYH11*)^47,59,60^. Consistent with this, C2 tumours are 4.8x (p = 1.78 x 10^-9^, Fisher’s Exact Test) more likely to be classified as ‘CAF-high’ than C1 tumours using a four-gene CAF index defined by Ko et al^48^. Indeed, three of the four CAF index genes (*TGFBI*, *TGFB2* and *FN1*) appear in the 938 DEG signature that separates C2 from C1 tumours (Supplementary Table S3).

## Discussion

In this study we hypothesized that by drawing upon several cervical cancer cohorts for which ‘omics data, clinical information and HPV typing were either available or for which we were able to profile samples ourselves, we would be able to gain further insight into CSCC – the most common histological cervical cancer subtype. Clustering of CSCCs according to the 10% most variable genes identified two clusters (C1 and C2) that bear resemblance to the keratin-high and keratin-low iClusters originally defined by TCGA^8^. Cluster membership is an independent predictor of 5-year OS and CSCCs can be accurately assigned to cluster using either a 938 gene expression signature or a 129 MVP DNA methylation signature, providing a means by which to gain prognostic information for cervical cancer patients. While HPV16 and the alpha-9 clade to which it belongs have been associated with longer PFS and OS in several studies^61–66^, the relationship between HPV genotype and cervical cancer prognosis remains unclear, as highlighted by a recent meta-analysis^7^. In our multivariate analyses, membership of the C2 cluster but not HPV type was an independent predictor of poor prognosis in both the discovery and validation cohorts and remained so when only HPV16-positive tumours in either cohort were considered. Possibly, the reason that HPV16 and other alpha-9 HPV types have been associated with more favourable outcomes in certain studies is that these viruses are more likely to cause C1-type tumours.

Adeno- and adenosquamous carcinomas, which are thought to arise from the columnar epithelium of the endocervix, have been linked to poor prognosis in cervical cancer ^3–6^ and to avoid differences due to histology, we focused our study entirely on CSCC. Interestingly, of the 14 keratin genes that are differentially expressed between C1 and C2 tumours, three (*KRT7*, *KRT8* and *KRT18*) that are upregulated in C2 were classified as marker genes for columnar-like tumours with a possible endocervical origin in a recent study that used single cell RNA-sequencing and lineage tracing experiments to explore cell-of-origin for CSCC and adenocarcinoma^67^. In contrast, C1 tumours display increased expression of *KRT5*, a marker of the squamous-like subtype with a proposed ectocervical origin identified by Chumduri et al^67^ (Fig. 3). Other signature genes (*TP63, CERS3, CSTA, CLCA2, DSC3* and *DSG3*) upregulated in C1 tumours are also markers of the squamous-like subtype, while further columnar-like marker genes (*MUC5B* and *RGL3*) are upregulated in C2 tumours (Supplementary Table S3). Squamous-like tumours are significantly enriched in the C1 sub-group, a C1 tumour is 4.9x more likely to be squamous-like than columnar-like or unclassified (Fisher Exact Test, p = 0.0003). This suggests that C2 tumours, although SCCs, harbour features associated with adenocarcinoma; possibly even hinting at a different cell-of-origin for C1 versus C2 tumours. The greater frequency with which alpha 7 HPV types are found in C2 SCCs is another feature shared with adenocarcinoma.

Our analysis suggests differences in the tumour immune microenvironment between C1 and C2 CSCCs, that are highly reproducible across cohorts from the USA, Europe and Uganda and that might explain the differential prognosis associated with these clusters. In addition to the high neutrophil:lymphocyte ratio, the increased expression of cytokines including IL-6, TGF-*β* and G-CSF and of the chemokines CXCL1-3 in C2 tumours suggests pro-tumourigenic (N2) polarisation of these neutrophils^68–73^, which is typical of tumours with a high NLR^74^. The observation that CSCCs occurring in HIV^+^ patients from the Ugandan/CGCI cohort are much more likely to be of the C2 subtype than those in HIV^-^ patients hints at a possible relationship between the immune competence of the patient and the likelihood of developing a C2 tumour. This requires further investigation but is consistent with greater evidence of existing anti-tumour immune responses in C1 tumours.

Finally, it is interesting to note that three targetable immune checkpoint proteins (B7-H3, NT5E and PD-L2) are expressed at higher levels in C2 tumours. In addition to its immune suppressive effects, B7-H3 has been linked to key processes that are upregulated in these tumours including EMT and angiogenesis, through the activation of NF-*κ*B signalling and the downregulation of E-cadherin expression^75,76^. Interestingly, the expression of B7-H3 and NT5E on CAFs has been linked to poor prognosis in gastric and colorectal cancer, respectively^40,77^. Also of relevance given our observation of differing CAF phenotype between clusters is the report that a CAF subtype (CAF-S1) identified in breast cancer that displays high levels of B7-H3 and NT5E expression is seen in tumours with low levels of CTL infiltration^78^. PD1/PD-L1 immune checkpoint blockade (pembrolizumab) was recently FDA-approved for first-line treatment of metastatic cervical cancer in combination with chemotherapy in patients whose tumours express PD-L1^79,80^, while CTLA4 blockade (Ipilimumab) has also shown promising activity, both as a single agent^81,82^ and in combination with PD1 blockade (Nivolumab)^83^. Efficacy of PD1 blockade in cervical cancer has been linked to the presence of a CD8+FoxP3+CD25+ T-cell subset^84^ and an important limitation of our study is the inability to differentiate between CD8+ T-cell phenotypes. Nonetheless, identification of alternative, targetable immune checkpoint molecules in C2 tumours provides a potential therapeutic strategy for a subset of cervical cancers that respond poorly to chemoradiotherapy and that, given their low overall levels of T-cell infiltrates, are maybe less likely to respond to PD1 blockade than C1 tumours.

In conclusion, we show that CSCCs can be categorised in two novel tumour types, C1 and C2, among which C1 tumours have a more favourable outcome. Although HPV16 is more likely to cause C1 tumours and HPV18 C2 tumours, HPV type is not an independent predictor of prognosis, suggesting it is the tumour type rather than the causative HPV type that is critical for the disease outcome. Notably, the key molecular and cellular characteristics of C1 and C2 tumours are consistent among cohorts from the US, Europe, and Sub-Saharan Africa. This suggests that the findings and underlying principle: that CSCC can develop along two trajectories associated with differing clinical behaviour that can be identified using defined gene expression or DNA methylation signatures, are of broad relevance.

## Methods

### Patient samples

All patients gave written, informed consent before inclusion. Samples from Bergen were collected in a population-based setting from patients treated at the Department of Obstetrics and Gynaecology, Haukeland University Hospital, Bergen, Norway, from May 2001 to May 2011. The study has been approved by the regional ethical committee (REK 2009/2315, 2014/1907 and 2018/591). For more details on sample collection see11,79. Samples from Innsbruck were collected and processed at the Department of Obstetrics and Gynaecology of the Medical University of Innsbruck. The study was reviewed and approved by the Ethics committee of the Medical University of Innsbruck (reference number: AN2016-0051 360/4.3; 374/5.4: ‘Biobank study: Validation of a DNA-methylation based signature in cervical cancer’) and conducted in accordance with the Declaration of Helsinki. Samples from Oslo (n = 268) were collected from patients participating in a previously published prospective clinical study80 approved by the Regional Committee for Medical Research Ethics in southern Norway (REK no. S-01129). Limited quantities of patient tumour samples and extracted DNA may remain and the distribution of these materials is subject to ethical approval at the institutions from which they were collected. Note that the cases in the Oslo cohort were not treated with surgery. The samples used for molecular analysis were diagnostic biopsies from the primary tumour. In all other cases, specimens were from resections of the primary tumour. Those interested in working with these samples should contact the authors to discuss their requirements.

### Dataset assembly

DNA methylation (Illumina Infinium 450k array) and RNAseq data were obtained for CESC from the TCGA data portal. TCGA mutation data were obtained from the MC3 project on SAGE Synapse (syn7214402). RNAseq data for the Uganda cohort was obtained from the TCGA data portal and DNA methylation (Illumina Infinium EPIC array) and mutation data from National Cancer Institute’s Genome Data Commons Publication Page at https://gdc.cancer.gov/about-data/publications/CGCI-HTMCP-CC-2020. DNA methylation (Illumina Infinium 450k array) and gene expression (Illumina HumanHT-12 V4.0 expression beadchip) data from the Oslo cohort were obtained from the Gene Expression Omnibus (GSE68339). RNAseq data were obtained for the Bergen cohort from dbGaP (phs000600/DS-CA-MDS ‘Genomic Sequencing of Cervical Cancers’) under the authorisation of project #14589 “Investigating the mechanisms by which viruses and carcinogens contribute to cancer development” and were converted to fastq files using SRA-dump from the SRA Toolkit (http://ncbi.github.io/sra-tools/). Kallisto81 was then used to quantify expression of GENCODE GrCh37 transcripts, repbase repeats and transcripts from 20 different high-risk HPV types with bias correction. Where IDAT files for 450k data were available, they were parsed using *minfi*82 and were subjected to Functional Normalisation83, followed by BMIQ-correction84 for probe type distribution (which was performed for all methylation data). For TCGA samples, viral type allocation was performed using VirusSeq85.

Only squamous cell carcinomas were considered in this study to avoid confounding from histology. Multidimensional visualisation of the molecular differences in histology was performed using Rtsne R package with parameters available in Supplementary Table S12, and the top 10% most variable genes using mean absolute deviation after pre filtering of low count genes (n = 1,385). Final cohort numbers and summaries are shown in Table 1.

### Generation of 450k methylation profiles

100ng DNA was bisulphite converted using the EZ DNA Methylation kit (Zymo Research) as per manufacturer’s instructions. Bisulphite converted DNA hybridised to the Infinium 450K Human Methylation array and processed in accordance with the manufacturer’s recommendations.

### HPV typing

HPV16 or 18 was detected in 208 samples from the Oslo cohort by PCR, using the primers listed in86. The PCR products were detected by polyacrylamide gene electrophoresis or the Agilent DNA 1000 kit (Agilent Technologies Inc, Germany). Samples from the Innsbruck cohort and the remaining non-HPV16/18 samples from the Oslo cohort (n=40) were HPV-typed by DDL Diagnostic Laboratory (Netherlands) using the SPF10 assay, in which a PCR-based detection of over 50 HPV types is followed by a genotyping assay (LIPA_25_) that identifies 25 HPV types (HPV 6, 11, 16, 18, 31, 33, 34, 35, 39, 40, 42, 43, 44, 45, 51, 52, 53, 54, 56, 58, 59, 66, 68/73, 70 and 74). If more than one HPV type was identified in a sample (e.g. HPV16 and HPV18), that sample was designated “Other” as HPV type in the study. HPV type data for the remaining samples were published previously^8,9,11^.

### Prognostic analyses and tumour clustering

Unsupervised consensus clustering was performed on TCGA SCC samples using r package ConsensusClusterPlus. After prefiltering of genes to remove those with low read counts (75% samples read count < 1), only the top 10% most variable genes using mean absolute deviation were considered for clustering (n = 1,385). 80% of tumours were sampled over 1000 iterations using all genes. PAM clustering algorithm was used and clustering distance was measured using Pearson’s correlation. An optimum number of clusters (K) of 2 was obtained by using the proportion of ambiguously clustered pairs (PAC) using thresholds of 0.1 and 0.9 to define the intermediate sub-interval. PAC was used as it accurately infers K87. Limma-voom on RNAseq data and limma on BMIQ and Functionally-normalised 450k and EPIC data were used to identify differentially expressed genes (DEGs, FDR = 0.01, FC > 2) and methylation variable positions (MVPs, FDR = 0.01, mean delta-Beta > 0.25) between the 2 clusters, C1 and C2. The 116 MVPs (Supplementary Table S13) common to the 450k and EPIC arrays were used to allocate clusters for the Ugandan cohort. The mean delta-Beta threshold for MVPs was determined as it delivered the highest concordance between DEG and MVP signature cluster allocation in the Bergen cohort (89.5%) and high concordance in the Ugandan cohort (91.5%). The caret R package and limma were used to develop an SVM using 5 iterations of 5-fold Cross-Validation using DEGs and MVPs to allocate RNAseq samples in Ugandan and Bergen cohorts, 450k samples in Bergen, Innsbruck and Oslo cohorts and EPIC samples in Ugandan cohort to these subgroups. Multidimensional visualisation using R package Rtsne was performed on the TCGA and European cohorts with available DNA methylation data combined using the 129 MVPs and parameters as shown in Supplementary Table S12.

Samples from our validation cohort, comprise of cases from three European centres (Bergen and Oslo in Norway and Innsbruck, Austria) and one African centre (Uganda) were binned into these categories, and were used for subsequent statistical analyses to identify genomic and microenvironmental correlates. Survival analyses of epigenetic allocations were carried out using Cox Proportional Hazards regression with age, tumour stage, HPV type, and with surgery, radiotherapy and chemotherapy (given/not given) as covariates. R packages used were survival and survminer. For all clinical analyses, stages were collapsed into Stages I, II, III and IV.

RNAseq data for Bergen and Ugandan samples, Illumina HumanWG-6 v3 microarray data for 137 of the Oslo samples and Illumina HumanHT-12 v4 microarray data for 109 of the Oslo samples were used to explore cluster allocation concordance accuracy between DEG and MVP signature cluster allocation. ROC curve and ssGSEA analysis were performed using R (scripts available at request).

#### Previous study comparison

140 TCGA samples from the core set analysis (TCGA, 2017) were present in our TCGA SCC cohort. Previous cluster analysis by TCGA (2017) and Chumduri *et al.* (2021) was compared with our C1 and C2 cluster allocation.

### Pathway analyses

Pathway and gene sets were analysed with Metascape^43^. Settings used were minimum gene set overlap of 10, p value cutoff of 0.01 and minimum enrichment of 1.5. All functional set, pathway, structural complex and miscellaneous gene sets were included in the analysis. Only hits with an FDR of less than 0.05 were included in final results.

### Mutational analyses

For TCGA data, mutation calls were obtained from SAGE synapse as called by the MC3 project. Mutations for the Bergen cohort were obtained from^11^. Ugandan mutation calls were obtained from National Cancer Institute’s Genome Data Commons Publication Page at https://gdc.cancer.gov/about-data/publications/CGCI-HTMCP-CC-2020. VCFs obtained for the Ugandan cohort samples were converted to maf files using R package vcf2maf, filtered for whole-exome mutations only, and combined. Significantly mutated genes (SMGs) were identified using dNdScv^16^ individually for the three cohorts. Hypermutated samples (>600 mutations^8^) were excluded from this analysis. A weighted approach was used to combine p values for each gene for the three cohorts. R package metapro^17^ function wFisher was used to perform this task. Genes were considered SMGs if after FDR correction of combined p values, q < 0.1. Analysis was repeated for only C1 and C2 samples individually. Two genes were removed from our list. *MUC4* was removed due to the large size of the gene and *GOLGA6L18* was removed as this gene and it’s aliases were not recognised by R package maftools88.

R package maftools was used to produce an oncoplot for SMGs, calculating tumour mutational burden for individual samples, SMG mutation frequency and mutational signatures for the combined cohorts. Binomial GLMs were used to estimate associations between C1 and C2 clusters and SMG mutation frequencies.

The estimated exposures of each sample to the identified mutational signatures were calculated using R package mutsignatures89 and converted to proportion of signature exposure per sample.

### Copy number analysis

450k total intensities (Methylated and Unmethylated values) were used to generate copy number profiles with normal blood samples from Renius et al90 as the germline reference. Functional normalisation83 was used to regress out technical variation across the reference and tumour datasets before merging and quantile normalisation was used to normalise combined intensities followed by Circular Binary Segmentation as previously described91. Median density peak correction was performed to ensure centering before further analysis. GISTIC2.092 was then used to identify regions of significant copy number change at both arm and gene levels. Candidate copy number changes were evaluated for association with cluster using binomial GLMs. The parameters chosen were a noise threshold of 0.1 with arm-level peel off and a confidence level of 0.95 was used to nominate genes targeted by copy number changes. Binomial regression was finally used to estimate rates of differential alteration.

### Reverse Phase Protein Assay analysis

Reverse Phase Protein Assay (RPPA) data for the core TCGA CESC samples were obtained from the NCI GDC Legacy Archive. Differentially expressed proteins between C1 and C2 clusters were determined using R package limma (FDR = 0.05, FC > 1.3).

### Tumour microenvironment analyses

MethylCIBERSORT^34^ was used to estimate tumour purity and abundances of nine other microenvironmental cellular fractions using TCGA and validation cohort methylation beta values. Fraction numbers were then normalised by cellular abundance and differences between clusters C1 and C2 were estimated using Wilcoxon’s rank sum test with Benjamini Hochberg correction for multiple testing. This analysis was performed separately on TCGA cohort and combined validation cohort, as well as on each individual cohort.

Cancer associated fibroblast associated gene set lists were obtained from Qian et al^56^. TCGA, Bergen and Ugandan cohort sample RNAseq data was combined and visualised for these gene set genes using R package NMF93.

### CAF Index calculation

For cohorts that RNAseq data was available (TCGA, Bergen and Uganda), a CAF index was calculated as described in Ko et al^48^. The median CAF index value was used as a threshold to allocate high or low CAF in tumour samples.

### Immunohistochemistry

Immunohistochemical staining of samples from the Innsbruck cohort was conducted by HSL-Advanced Diagnostics (London, UK) using the Leica Bond III platform with Leica Bond Polymer Refine detection as per manufacturer’s recommendations. Sections from a series of 17 tumour samples from the validation cohort were stained for CD8 (mouse monoclonal 4B11, Leica Biosystems PA0183, used as supplied for 15 minutes at room temperature. HIER was performed on-board using Leica ER2 solution (high pH) for 20 minutes), CD68 (mouse monoclonal PGM1, Agilent M087601-2, used at a dilution of 1/50 for 15mins at room temperature. HIER was performed on-board using Leica ER1 solution (low pH) for 20 minutes) or MPO (rabbit polyclonal, Agilent A039829-2, used at a dilution of 1/4000 for 15 minutes at room temperature without epitope retrieval. Scoring was performed blinded to cluster membership by a histopathologist (JM) as follows: 0 = no positive cells / field (200X magnification); 1 = 1 – 10 positive cells; 2 = 11 – 100 positive cells; 3 = 101 – 200 positive cells; 4 = 201 = 300 positive cells; 5 = over 300 positive cells.

For the Oslo cohort, manual CD8 staining was conducted using the Dako EnVision^TM^ Flex+ System (K8012, Dako). Deparaffinization and unmasking of epitopes were performed using PT-Link (Dako) and EnVision^TM^ Flex target retrieval solution at a high pH. The sections were incubated with CD8 mouse monoclonal antibody (clone 4B11, 1:150, 0.2 µg IgG_2b_/ml) from Novocastra (Leica Microsystems, Newcastle Upon Tyne, UK) for 45 minutes. All CD8 series included positive controls. Negative controls included substitution of the monoclonal antibody with mouse myeloma protein of the same subclass and concentration as the monoclonal antibody. All controls gave satisfactory results. CD8 pathology scores were given to each sample (blinded to cluster membership) for connective tissue only, tumour only and both as follows: 0 = no positive: 1 = <10% CD8 positive cells; 2 = 10-25% CD8 positive cells; 3 = 25-50% CD8 positive cells; 4 = >50% CD8 positive cells. For digital quantification scanned images of all sections at a high resolution of 0.46 um/pixel (20x), which was reduced to 0.92 um/pixel for analysis, were used. Digital score was calculated by quantifying the area fraction of stained CD8 cells in relation to the entire section in the digital assessment.

## Data availability

Illumina Infinium 450k array DNA methylation data generated in-house from Bergen and Innsbruck validation cohort samples have been deposited in the Gene Expression Omnibus (accession number GSEXXXXX (to be deposited upon publication)). For detailed information on all other datasets see ‘Dataset Assembly’.

## Code availability

All packages used have been published, are freely available and are referenced in the methods. R markdowns used to run the analyses specific to this study are available from the authors on request.

## Acknowledgements

AC was supported by postgraduate research scholarships from UCL and received additional research support from a Debbie Fund grant to KC and TRF. TRF was supported by Rosetrees Trust (M229-CD1), Cancer Research UK (A25825), the Biotechnology and Biosciences Research Council (Grant Ref: BB/V010271/1), the Royal Society (IEC\R2\202256) and the Global Challenges Doctoral Centre at the University of Kent. DNA methylation data were generated through funding provided by the Debbie Fund and the results shown here are in part based upon data generated by the TCGA Research Network: https://www.cancer.gov/tcga and the Cancer Genome Characterization Initiative: https://ocg.cancer.gov/programs/cgci. AF was supported by grants from the MRC (MR/M025411/1), PCUK(MA-TR15-009), BBSRC (BB/R009295/1), TUF, Orchid and the UCLH BRC. The authors dedicate this manuscript to the late Dr Helga Salvesen, a wonderful collaborator and colleague who played a key role in the project.

## Supplementary Figure Legends

**Supplementary Figure S1 – tSNE clustering by histology in cervical cancer cohorts.** Unsupervised tSNE analysis using top 10% most variable genes for cervical cancer cohorts **a)** TCGA (1385 most variable genes), **b)** Ugandan (1371) and **c)** Bergen (1430). Concordance of most variable genes was high amongst the 3 cohorts **(d)**.

**Supplementary Figure S2 Consensus clustering using ConsensusClusterPlus. a)** Consensus CDF plot. PAC score = CDF at 0.9 consensus index – CDF at 0.1 consensus index for each curve. **b)** Delta area plot used in decision of optimum number of clusters.

**Supplementary Figure S3 Genes that are both differentially expressed and differentially methylated between C1 and C2 subgroups.** Datapoints represent methylated variable positions (in either the 3’UTR, body of gene, intergenic region or gene promoter) in genes that are also differentially expressed between C1 and C2 subgroups. Datapoints in the top left quadrant are MVPs that are hypomethylated in genes that are also upregulated in C2 tumours. Those in the bottom right quadrant are hypermethylated in genes that are downregulated in C2 tumours.

**Supplementary Figure S4 Concordance between gene expression and DNA methylation-derived cluster membership. a)** The percentage of samples that are designated the same cluster allocation by gene expression signature and methylation signatures based on varying delta Beta thresholds. **b)** ROC curves showing the accuracy with which C1 or C2 cluster membership can be predicted using DNA methylation differences (MVPs) in samples from the validation cohorts for which either RNA-seq (Bergen, n=37, and Uganda, n=94, HPV+ SCC cases), Illumina HumanHT-12 V4.0 expression beadchip array (Oslo SCC cases, n=109) or Illumina HumanWG-6 v3.0 expression beadchip array (Oslo SCC cases, n=139) gene expression data were available. **c)** Single sample gene set enrichment analysis (ssGSEA) for validation cohorts used in panel B. The y-axis represents the ssGSEA score for each sample, compared with the genes from the C2 gene expression signature. P-values from Wilcoxon rank-sum test.

**Supplementary Figure S5 Validation SCC cohorts. a)** Ugandan validation cohort clustering based on 116 MVP signature. Kaplan-meier curves for **b)** HPV16+ European validation cohort SCC patients; **c)** European validation cohort SCC patients without chemotherapy treatment and **d)** 5 year survival for the 5 individual cohorts in this study.

**Supplementary Figure S6 Elevation of epithelial mesenchymal transition (EMT) score is evident in C2 tumours. a)** EMT score derived by TCGA for 140 HPV+ squamous TCGA cervical cancer tumours in our study. EMT score is higher in C2 tumours.

**Supplementary Figure S7 Mutational signatures of combined HPV+ squamous cervical cancer cohorts.** COSMIC mutational signatures identified in combined HPV+ squamous cervical cancer cohort including genomic data from TCGA, Bergen and Ugandan cohorts.

**Supplementary Figure S8 Increased levels of YAP in tumours with YAP1 amplification.** YAP1 expression **(a)**, and YAP protein levels **(b)** unphosphorylated, **c)** phosphorylated) are higher in tumours that contain YAP1 amplifications.

**Supplementary Figure S9 Differences in immune microenvironment between SCC subgroups in individual cohorts.** Median abundances (x-axis) and median differences (%, y-axis) for different cell types estimated using MethylCIBERSORT, with significant differences in orange for cohorts from **a)** Bergen, **b)** Innsbruck, **c)** Oslo and **d)** Uganda. C2 tumours display increased neutrophil:CTL ratios as estimated using MethylCIBERSORT for **e)** TCGA discovery cohort and **f)** combined validation cohorts.

**Supplementary Figure S10 Immune cell ratios by cluster using MethylCIBERSORT estimates. a)** Neutrophil:CD19 estimate ratios for combined cohorts. **b)** Neutrophil:Treg estimate ratios for combined cohorts.

**Supplementary Figure S11 Comparison of MethylCIBERSORT estimates and immunohistochemistry(IHC)-based scoring.** Correlations between MethylCIBERSORT estimates and IHC-based scoring for **a)** CD8+ T-cells, **b)** neutrophils (MPO+), **c)** CD8+ T-cell:neutrophil ratio in 14 SCCs from the Innsbruck validation cohort and **d)** CD8+ T-cells for 229 SCCs from the Oslo validation cohort. Trendlines are derived from linear modelling, shaded areas represent 95% CI of trendlines.

**Supplementary Figure S12 Upregulation of immune checkpoint genes in C2 SCCs.** Upregulation of **a)** *B7-H3* (*CD276*), **b)** *NT5E* (*CD73*) and **c)** *PD-L2* (*PDCD1LG2*) was observed in poor prognosis C2 tumours. Analysis performed with RNA-seq data from TCGA, Bergen and Ugandan cohorts.

## Supplementary Tables

Table S1 - Clinical and pathaologic characteristics of TCGA squamous cervical cancer cohort samples

Table S2 - Top 10% most variable genes in TCGA squamous cervical cancer cohort

Table S3 - 938 Differentially expressed genes between TCGA squamous cervical cancer clusters C1 and C2

Table S4 - 5 year survival uni- and multivariate analysis for HPV16+ patients in squamous cervical cancer cohorts

Table S5 - 129 MVP signature probes (European validation cohorts)

Table S6 - Combined validation cohort cluster allocation

Table S7 - Breakdown of tumour stage in C1 and C2 cluster by percentage

Table S8 - Clusters and EMT scores for TCGA squamous cervical cancer samples

Table S9 - Significantly mutated genes using dNdSCV analysis and combining cohorts

Table S10 - Mutation frequency in SMGs observed in previous studies

Table 11 - Gene set enrichment analysis of C2 gene expression signature genes using Metascape

Table S12 - Paramaters for TSNE multidimensional visualisation analyses

Table S13 - 116 MVP signature probes (Ugandan validation cohort)

