## Supplementary Fig. S1 for "Integrated analysis of cervical squamous cell carcinoma cohorts from three continents reveals conserved subtypes of prognostic significance"

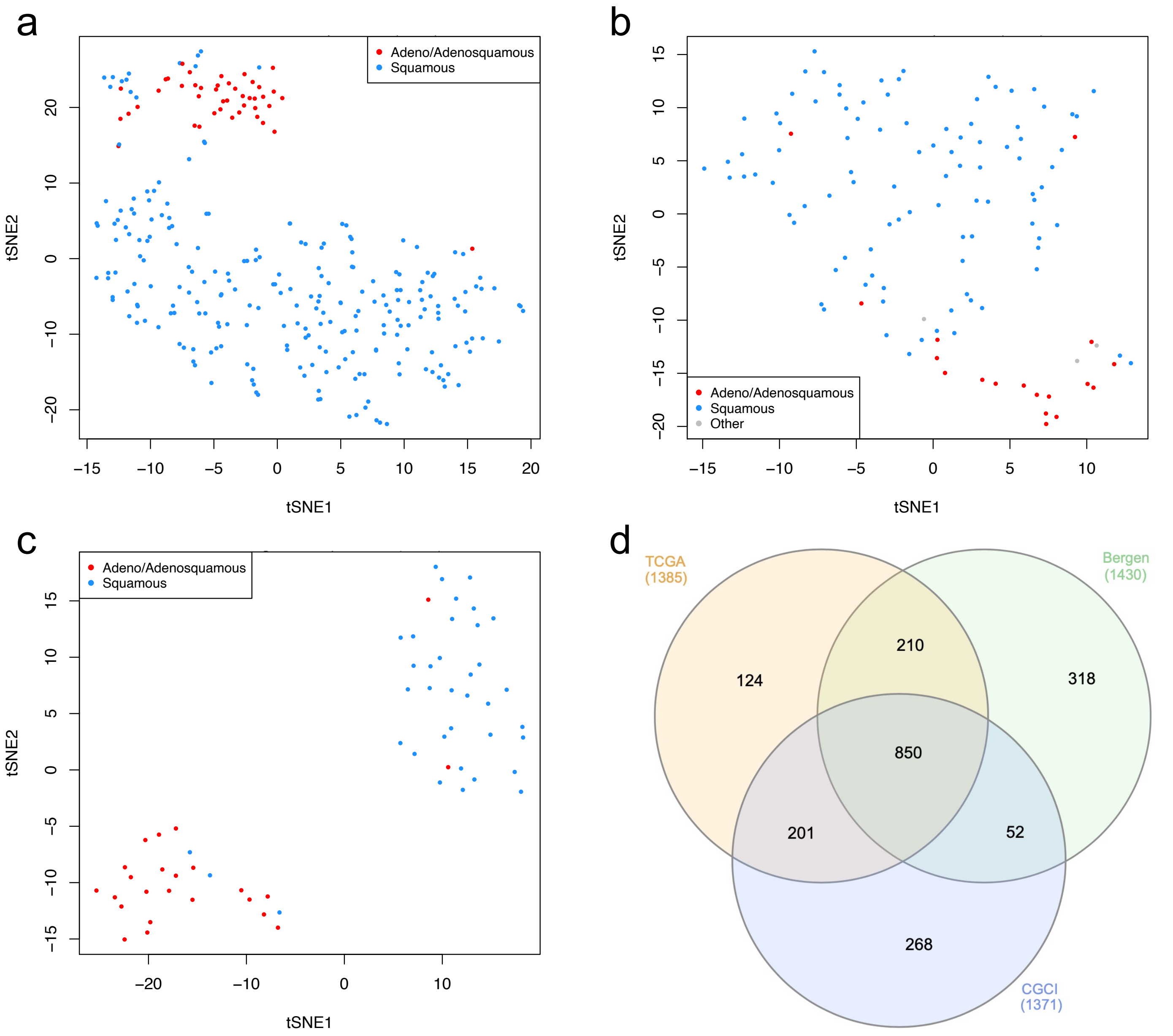


**Supplementary Figure S1 – tSNE clustering by histology in cervical cancer cohorts.** Unsupervised tSNE analysis using top 10% most variable genes for cervical cancer cohorts **a)** TCGA (1385 most variable genes), **b)** Ugandan (1371) and **c)** Bergen (1430). Concordance of most variable genes was high amongst the 3 cohorts **(d)**.
