## Supplementary Fig. S2 for "Integrated analysis of cervical squamous cell carcinoma cohorts from three continents reveals conserved subtypes of prognostic significance"

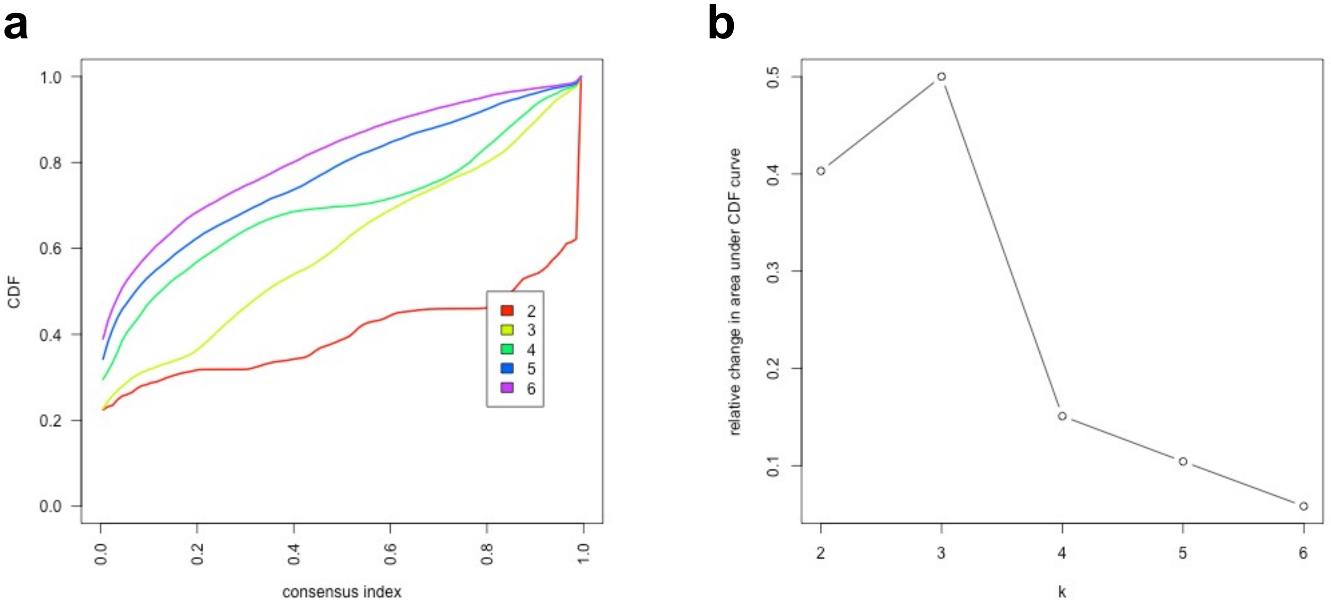


**Supplementary Figure S2 Consensus clustering using ConsensusClusterPlus. a)** Consensus CDF plot. PAC score = CDF at 0.9 consensus index – CDF at 0.1 consensus index for each curve. **b)** Delta area plot used in decision of optimum number of clusters.
