## Supplementary Fig. S3 for "Integrated analysis of cervical squamous cell carcinoma cohorts from three continents reveals conserved subtypes of prognostic significance"

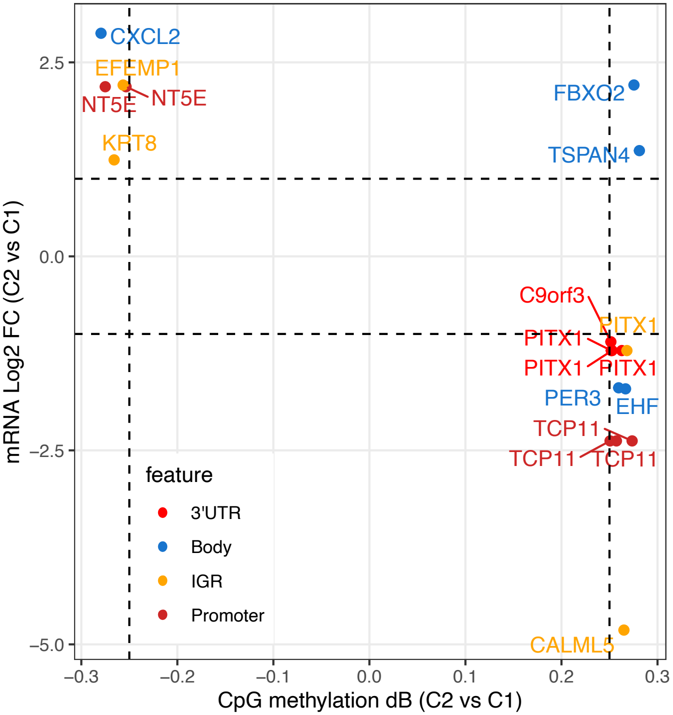


**Supplementary Figure S3 Genes that are both differentially expressed and differentially methylated between C1 and C2 subgroups**. Datapoints represent methylated variable positions (in either the 3'UTR, body of gene, intergenic region or gene promoter) in genes that are also differentially expressed between C1 and C2 subgroups. Datapoints in the top left quadrant are MVPs that are hypomethylated in genes that are also upregulated in C2 tumours. Those in the bottom right quadrant are hypermethylated in genes that are downregulated in C2 tumours.
