## Supplementary Fig. S4 for "Integrated analysis of cervical squamous cell carcinoma cohorts from three continents reveals conserved subtypes of prognostic significance"

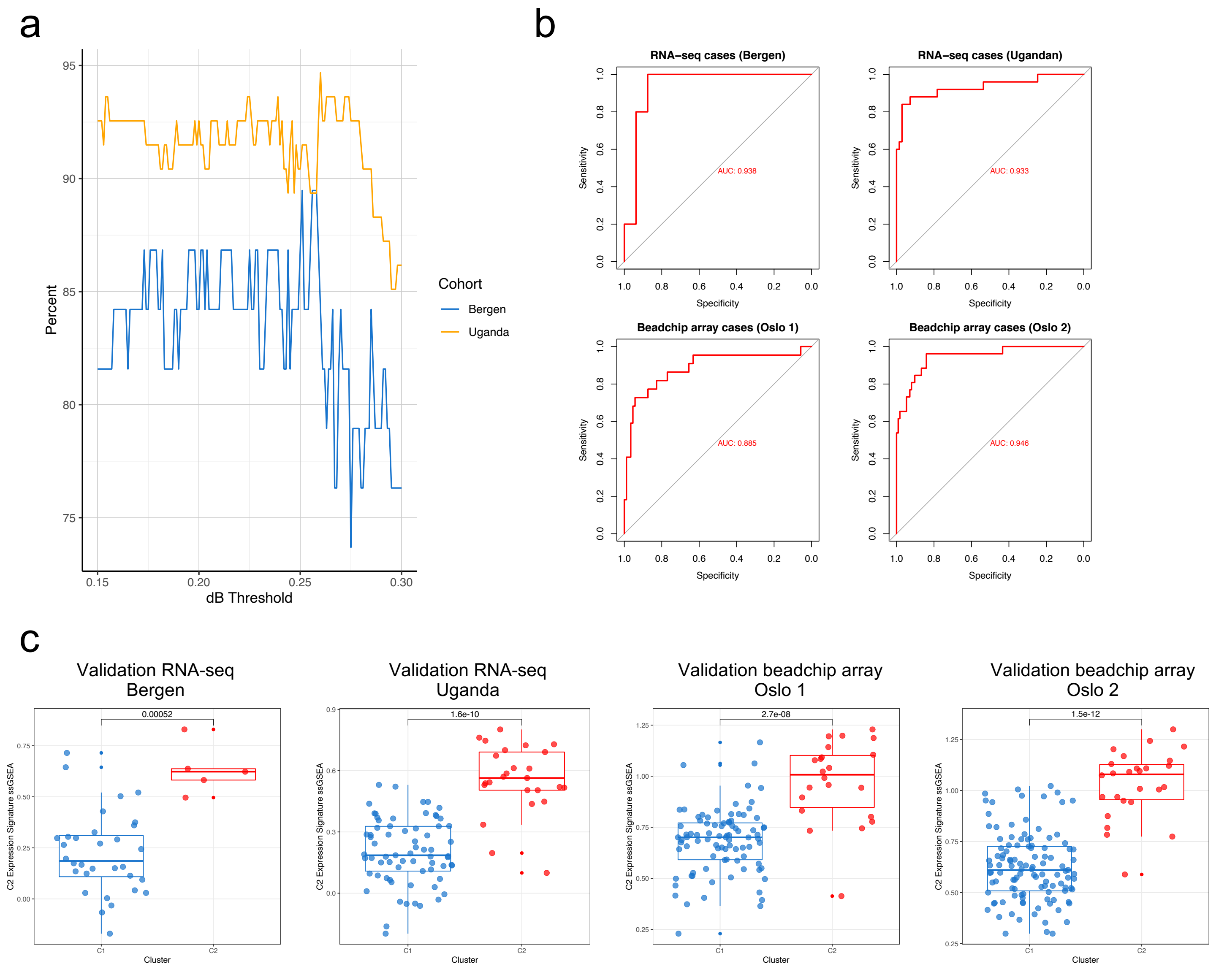


**Supplementary Figure S4 Concordonce between gene expression and DNA methylation-derived cluster membership. a)** The percentage of samples that are designated the same cluster allocation by gene expression signature and methylation signatures based on varying delta Beta thresholds. **b)** ROC curves showing the accuracy with which C1 or C2 cluster membership can be predicted using DNA methylation differences (MVPs) in samples from the validation cohorts for which either RNA-seq (Bergen, n=37, and Uganda, n=94, HPV+ SCC cases), Illumina HumanHT-12 V4.0 expression beadchip array (Oslo SCC cases, n=109) or Illumina HumanWG-6 v3.0 expression beadchip array (Oslo SCC cases, n=139) gene expression data were available. **c)** Single sample gene set enrichment analysis (ssGSEA) for validation cohorts used in panel B. The y-axis represents the ssGSEA score for each sample, compared with the genes from the C2 gene expression signature. P-values from Wilcoxon rank-sum test.
