## Supplementary Fig. S5 for "Integrated analysis of cervical squamous cell carcinoma cohorts from three continents reveals conserved subtypes of prognostic significance"

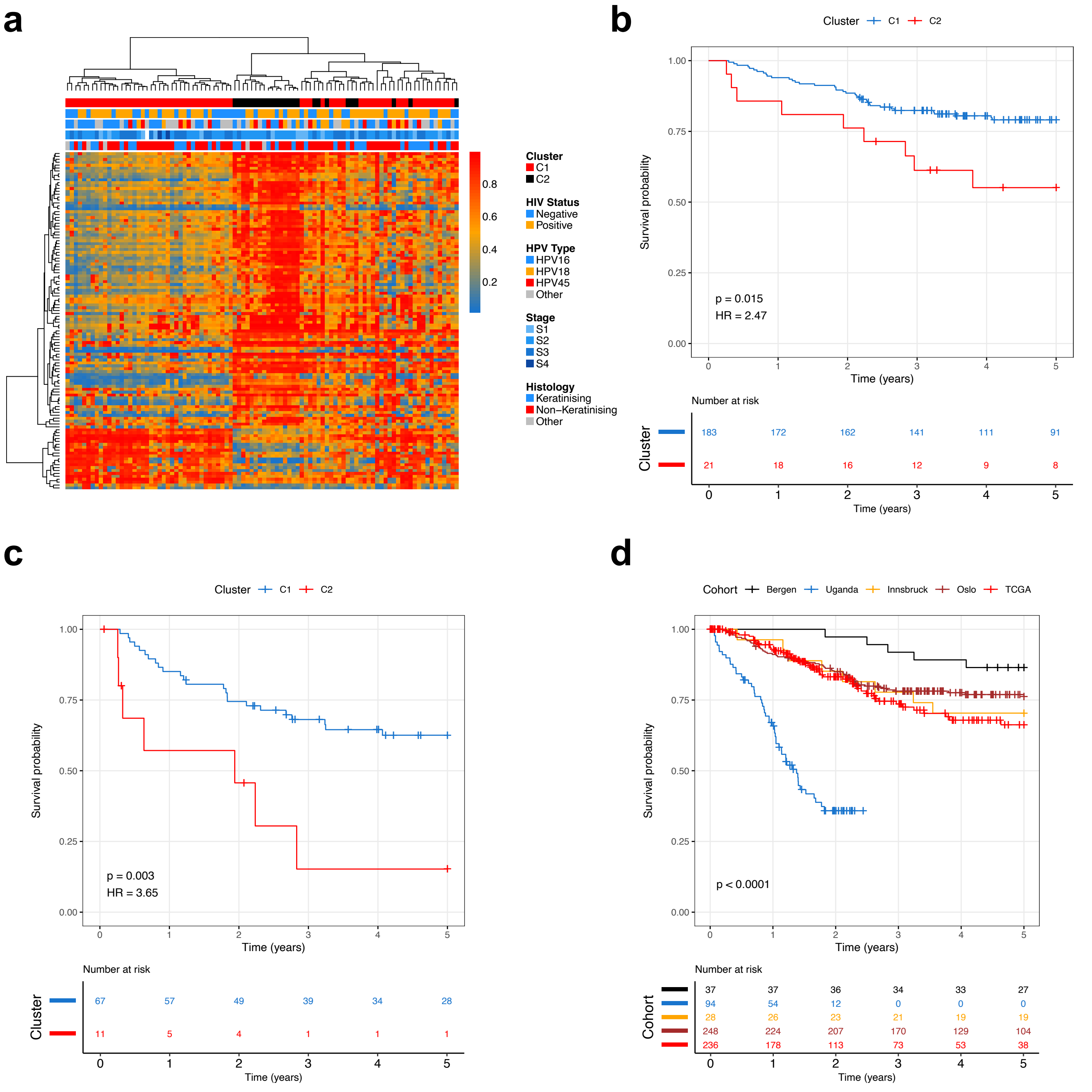


**Supplementary Figure S5 Validation SCC cohorts. a)** Ugandan validation cohort clustering based on 116 MVP signature. Kaplan-meier curves for **b)** HPV16+ European validation cohort SCC patients; **c)** European validation cohort SCC patients without chemotherapy treatment and **d)** 5 year survival for the 5 individual cohorts in this study.
