## Supplementary Fig. S6 for "Integrated analysis of cervical squamous cell carcinoma cohorts from three continents reveals conserved subtypes of prognostic significance"

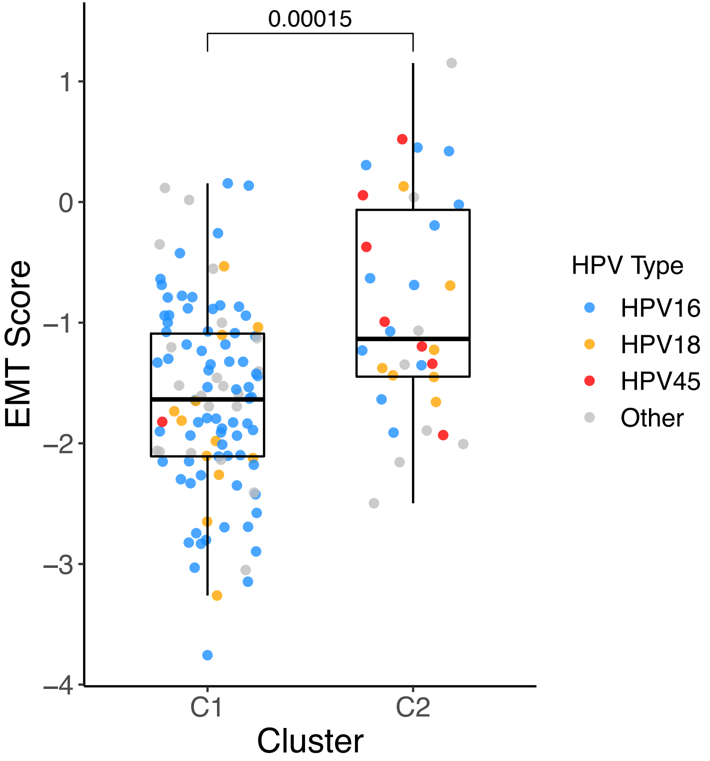


**Supplementary Figure S6 Elevation of epithelial mesenchymal transition (EMT) score is evident in C2 tumours.** EMT score derived by TCGA for 140 HPV+ squamous TCGA cervical cancer tumours in our study. EMT score is higher in C2 tumours.
