## Supplementary Fig. S7 for "Integrated analysis of cervical squamous cell carcinoma cohorts from three continents reveals conserved subtypes of prognostic significance"

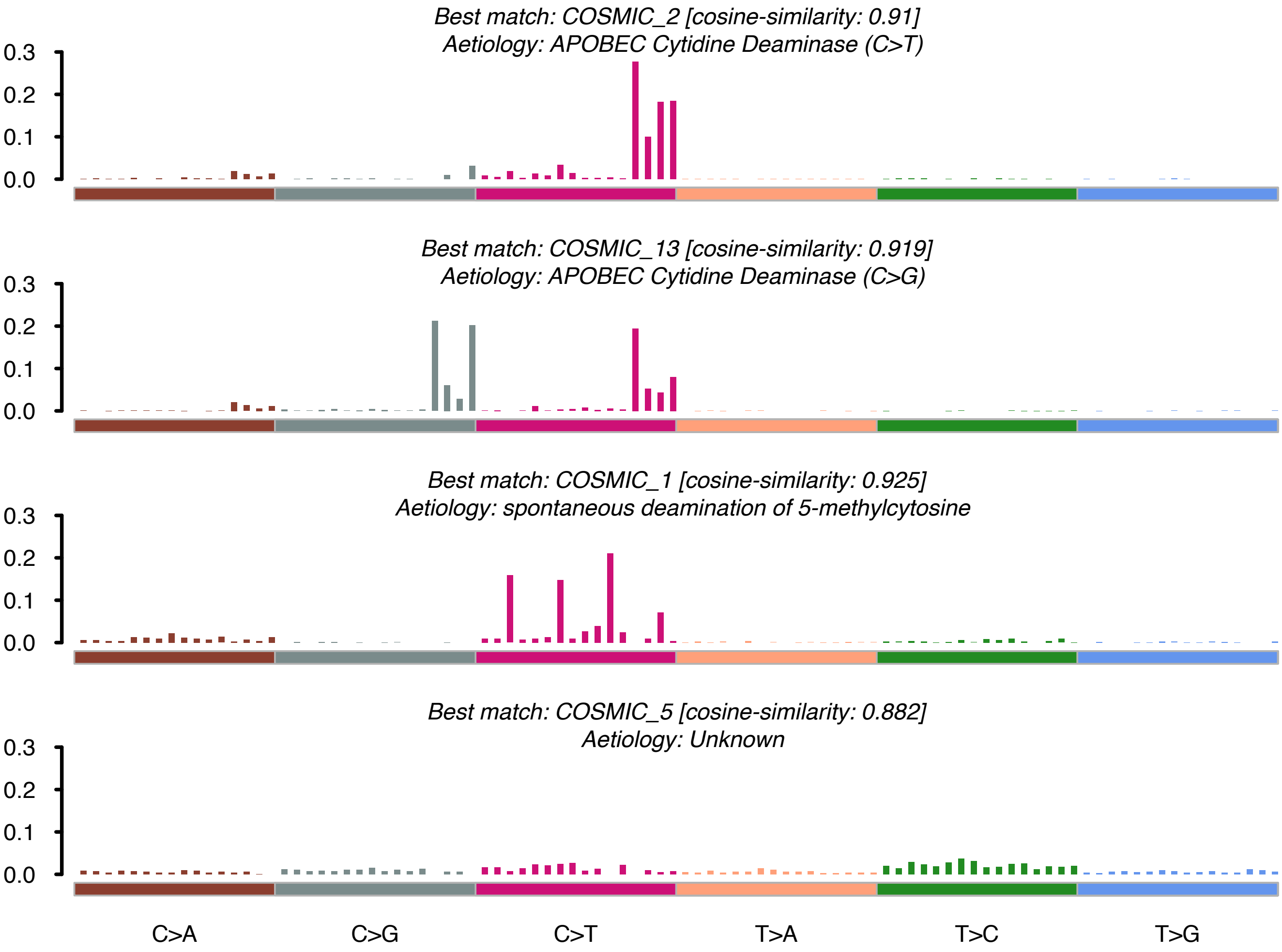


**Supplementary Figure S7 Mutational signatures of combined HPV+ squamous cervical cancer cohorts.** COSMIC mutational signatures identified in combined HPV+ squamous cervical cancer cohort including genomic data from TCGA, Bergen and Ugandan cohorts
