## Supplementary Fig. S8 for "Integrated analysis of cervical squamous cell carcinoma cohorts from three continents reveals conserved subtypes of prognostic significance"

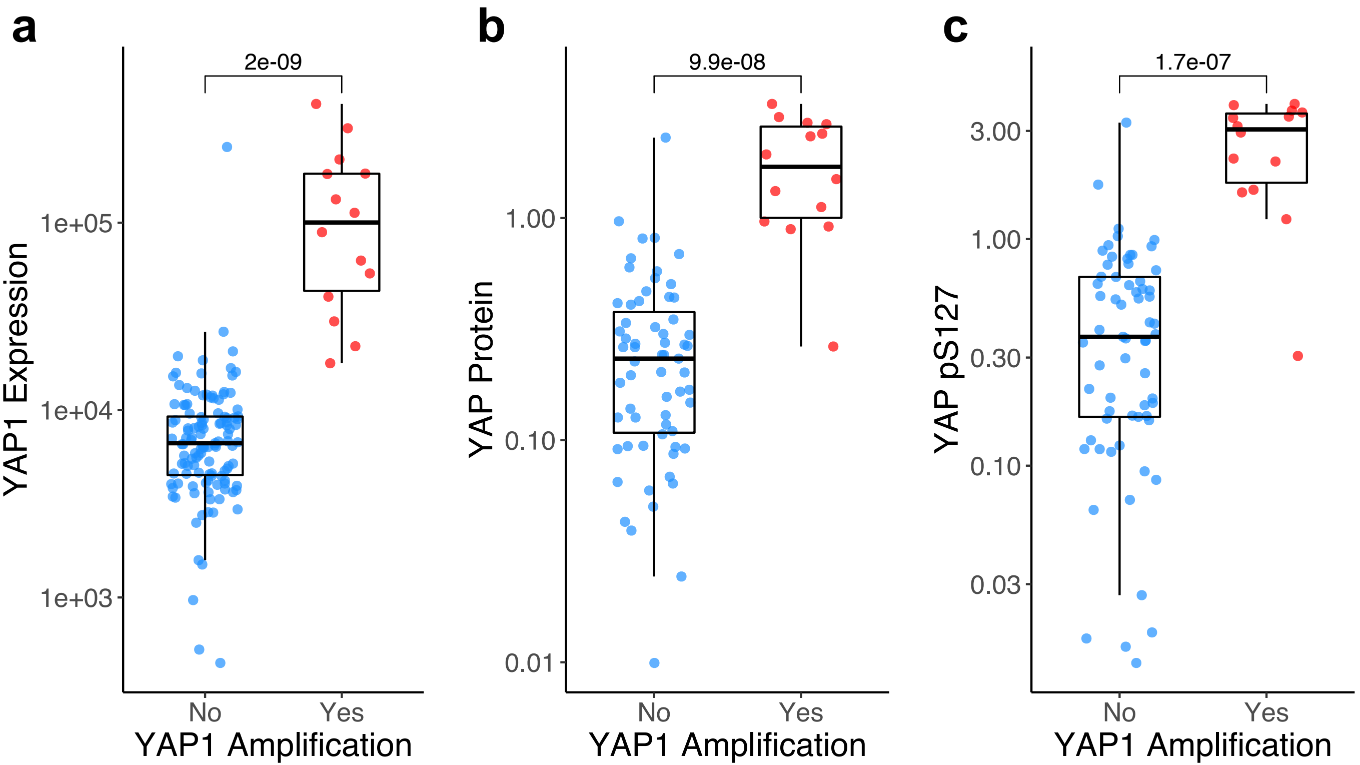


**Supplementary Figure S8 Increased levels of YAP in tumours with YAP1 amplification.** YAP1 expression **(a)**, and YAP protein levels **(b)** unphosphorylated, **c)** phosphorylated) are higher in tumours that contain YAP1 amplifications.
