## Supplementary Fig. S9 for "Integrated analysis of cervical squamous cell carcinoma cohorts from three continents reveals conserved subtypes of prognostic significance"

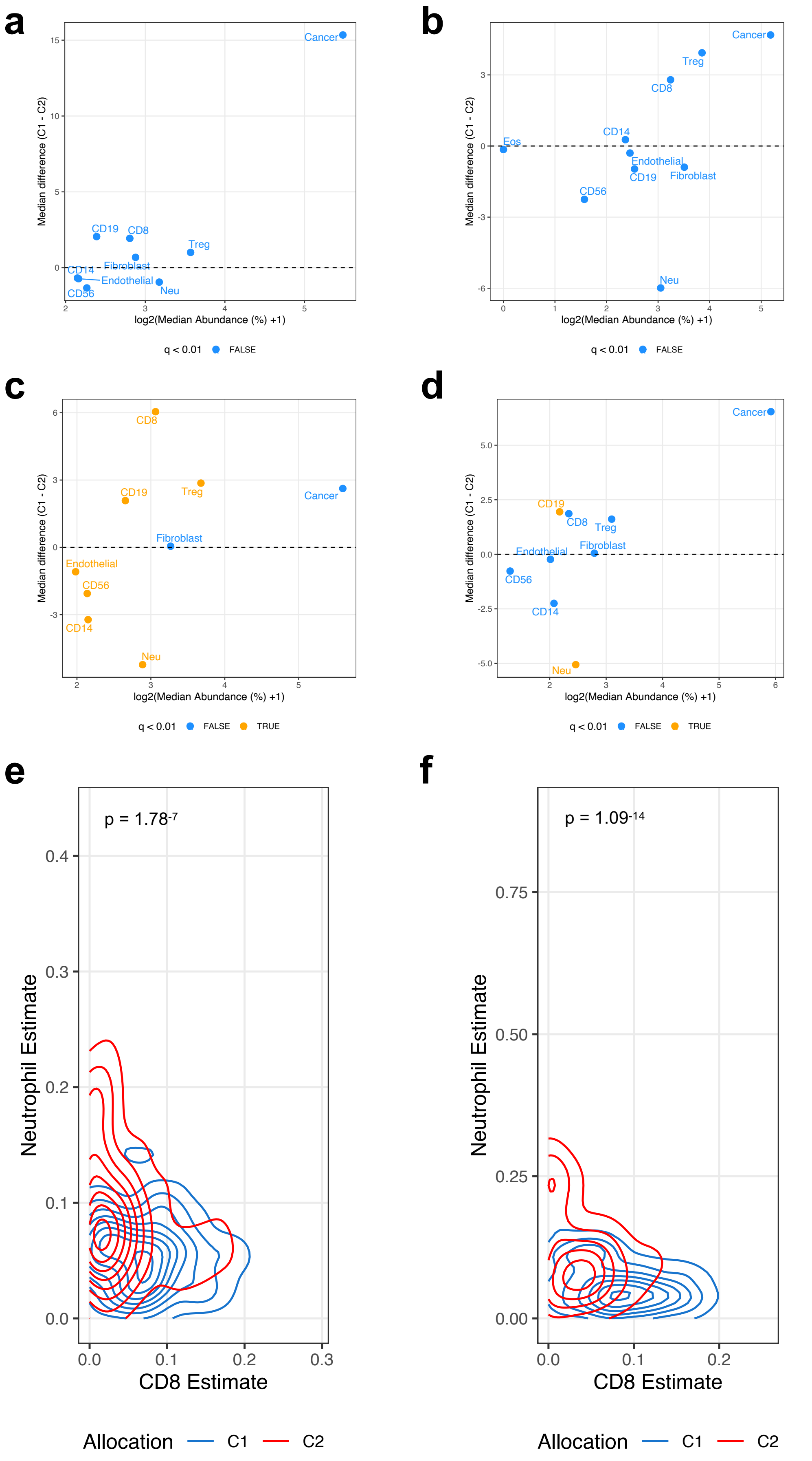


**Supplementary Figure S9 Differences in immune microenvironment between SCC subgroups in individual cohorts.** Median abundances (x-axis) and median differences (%, y-axis) for different cell types estimated using MethylCIBERSORT, with significant differences in orange for cohorts from **a)** Bergen, **b)** Innsbruck, **c)** Oslo and **d)** Uganda. C2 tumours display increased neutrophil:CTL ratios as estimated using MethylCIBERSORT for **e)** TCGA discovery cohort and **f)** combined validation cohorts.
