## Supplementary Fig. S10 for "Integrated analysis of cervical squamous cell carcinoma cohorts from three continents reveals conserved subtypes of prognostic significance"

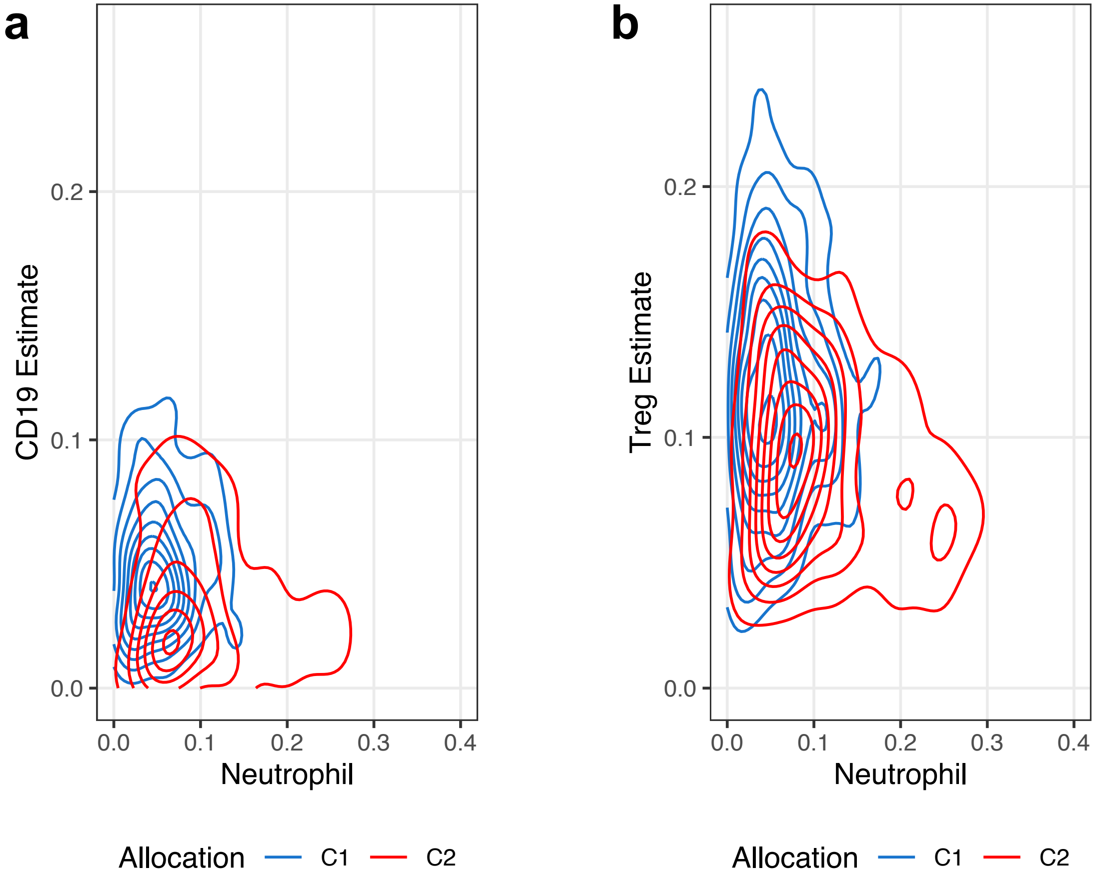


**Supplementary Figure S10 Immune cell ratios by cluster using MethylCIBERSORT estimates. a)** Neutrophil:CD19 estimate ratios for combined cohorts. **b)** Neutrophil:Treg estimate ratios for combined cohorts.
