## Supplementary Fig. S11 for "Integrated analysis of cervical squamous cell carcinoma cohorts from three continents reveals conserved subtypes of prognostic significance"

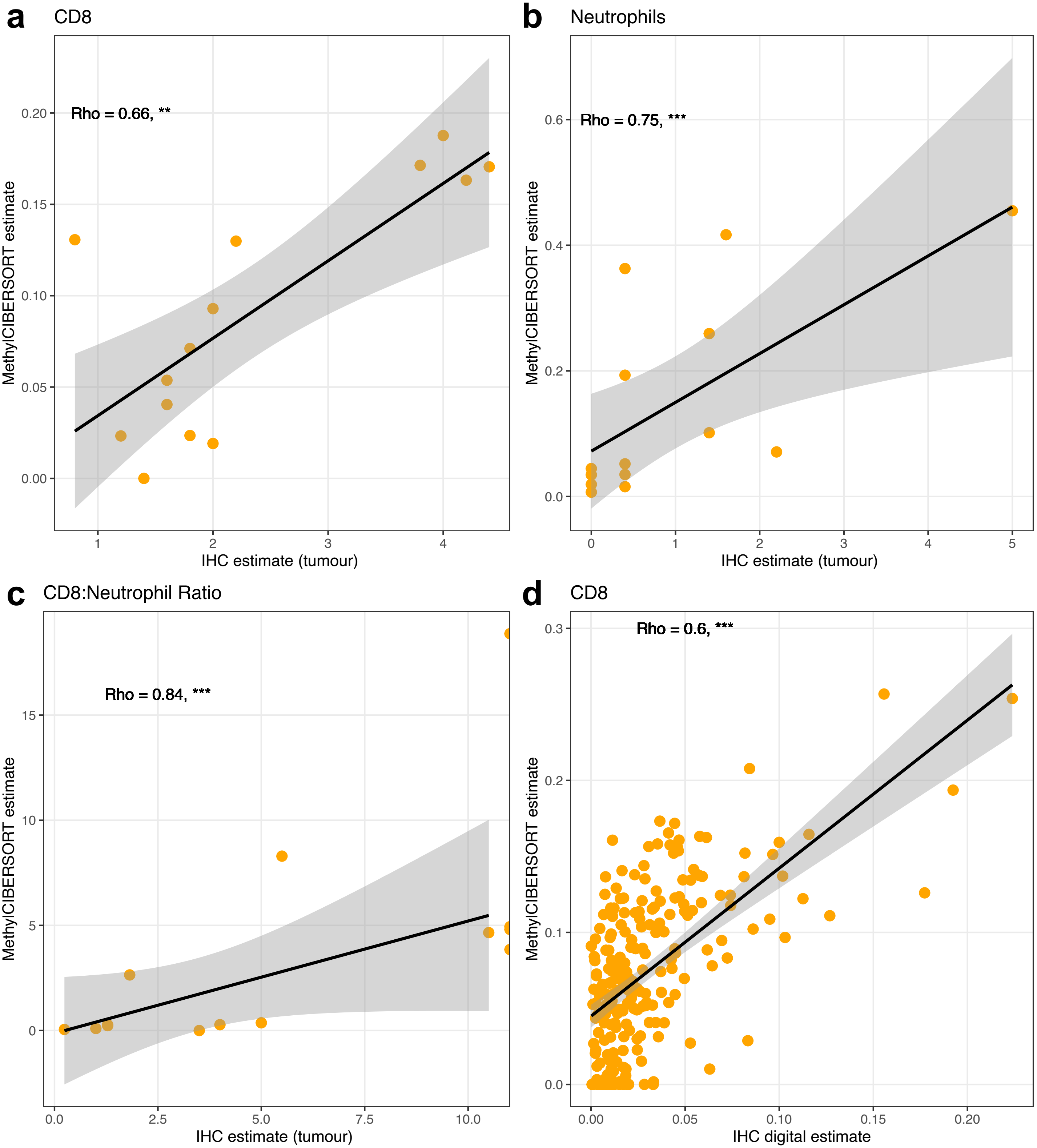


**Supplementary Figure S11 Comparison of MethylCIBERSORT estimates and immunohistochemistry(IHC)-based scoring.** Correlations between MethylCIBERSORT estimates and IHC-based scoring for **a)** CD8+ T-cells, **b)** neutrophils (MPO+), **c)** CD8+ T-cell:neutrophil ratio in 14 SCCs from the Innsbruck validation cohort and **d)** CD8+ T-cells for 229 SCCs from the Oslo validation cohort. Trendlines are derived from linear modelling, shaded areas represent 95% CI of trendlines.
