## Supplementary Fig. S12 for "Integrated analysis of cervical squamous cell carcinoma cohorts from three continents reveals conserved subtypes of prognostic significance"

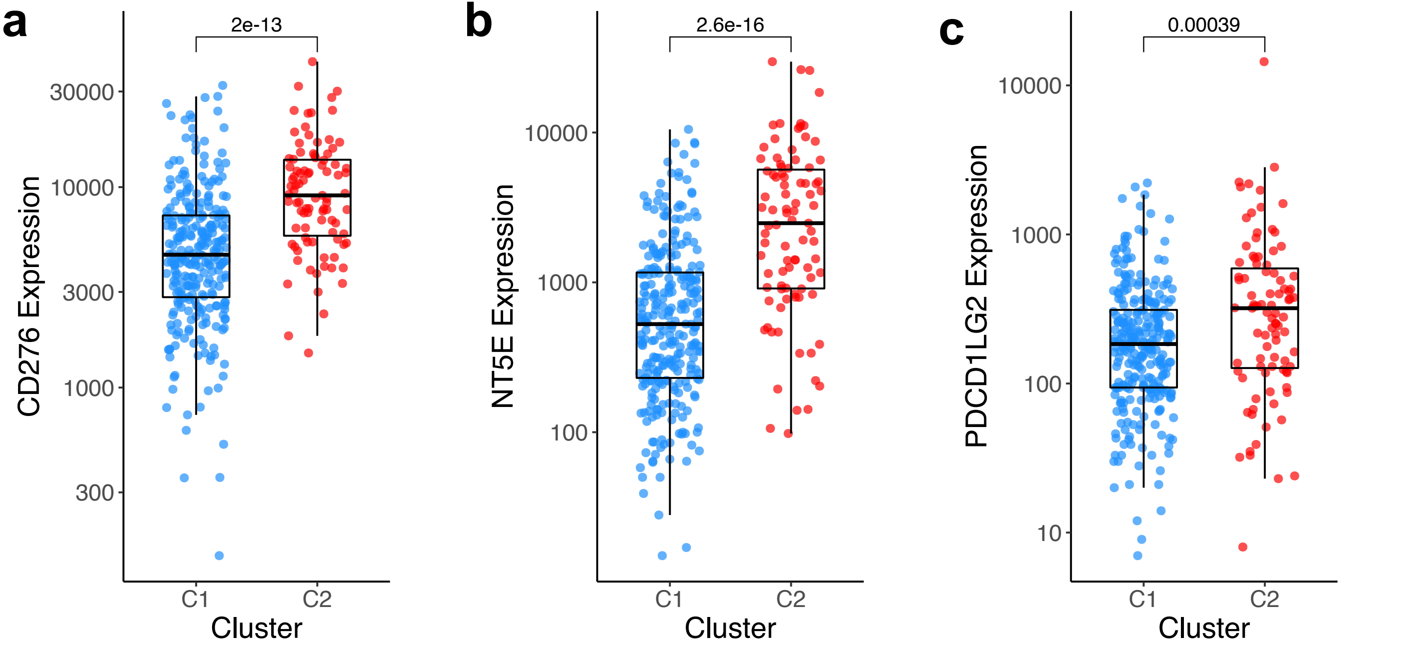


**Supplementary Figure S12 Upregulation of immune checkpoint genes in C2 SCCs.** Upregulation of **a)** *B7-H3* (*CD276*), **b)** *NT5E* (*CD73*) and **c)** *PD-L2* (*PDCD1LG2*) was observed in poor prognosis C2 tumours. Analysis performed with RNA-seq data from TCGA, Bergen and Ugandan cohorts.
